# A platform for high-throughput and ultrasensitive immunopeptidomics

**DOI:** 10.64898/2026.02.23.707388

**Authors:** Adillah Gul, Laura Van Moortel, Patrick Willems, Ilke Aernout, Laura Pedró-Cos, Kia C. Ferrell, Katie Boucher, An Staes, Simon Devos, Ine Lentacker, Bart Vandekerckhove, Caroline Demangel, Fabien Thery, Francis Impens

**Affiliations:** VIB Center for Medical Biotechnology, VIB, Ghent, Belgium; Department of Biomolecular Medicine, Ghent University, Ghent, Belgium; Ghent Research Group on Nanomedicines, Ghent University, Ghent, Belgium; Institut Pasteur, Université Paris Cité, Inserm U1224, Immunobiology and Therapy Unit, Paris, France; VIB Proteomics Core, VIB, Ghent, Belgium; Department of Diagnostic Sciences, Ghent University, Ghent, Belgium; Cancer Research Institute Ghent (CRIG), Ghent, Belgium; GMP Unit cell Therapy, Ghent University Hospital, Ghent, Belgium

**Author notes:** Correspondence to: Francis Impens and Fabien Thery VIB-UGent Center for Medical Biotechnology Technologiepark-Zwijnaarde 75 9052 Gent Belgium. These authors contributed equally (Adillah Gul, Laura Van Moortel, Patrick Willems).

## Abstract

Mass spectrometry (MS)-based immunopeptidomics is a powerful approach for untargeted discovery of peptides presented on major histocompatibility complex (MHC) molecules, which can guide the selection of vaccine antigens and immunotherapy targets. First-generation immunopeptidomics workflows require processing of hundreds of millions of cells using lengthy, manual procedures. More recent approaches focus on increasing either sensitivity or throughput, but rarely combine both aspects. Here, we describe a semi-automated immunopeptidomics platform that combines high sensitivity with high throughput by implementing highly optimized conditions for immunoprecipitation, elution and purification of MHC class I and II peptides on a 96-well positive-pressure device. Upon analysis of 25% of the eluate from 16 million cells, our workflow identified over 13,500 MHC I and 6,000 MHC II peptides on a timsTOF SCP mass spectrometer, operating in DDA-PASEF mode. Exploring the sensitivity limits of our platform, we identified over 1,000 MHC I peptides from as few as 20,000 JY cells. Validating the platform’s performance for quantitative biological discovery, we report the identification of known and novel bacterial immunopeptides from U937 macrophages infected with *Listeria monocytogenes* or Bacillus Calmette-Guérin (BCG). Together, our optimized immunopeptidomics platform enables robust immunopeptide detection from lower-input samples in a high-throughput fashion, enabling its use for biological applications where sample amounts are limiting.

## INTRODUCTION

The immunopeptidome is the repertoire of peptides presented by major histocompatibility complex (MHC) molecules, also known as human leukocyte antigens (HLAs) in humans, on the surface of cells (1). These MHC-bound peptides, called immunopeptides, are derived from self and non-self proteins processed inside the cell (2). Human MHC class I molecules present peptides that are 8- to 12-amino acids in length that result from proteasomal degradation of intracellular proteins. MHC class II molecules typically present 13- to 18-mer peptides generated by the uptake and lysosomal degradation of extracellular proteins. Peptides presented on MHC-I and MHC-II are recognized by CD8^+^ and CD4^+^ T cells, respectively. Therefore, the immunopeptidome plays a critical role in adaptive immunity, facilitating recognition of peptides derived from pathogens, tumor antigens, and aberrantly expressed self-proteins by T cells (3–6). Over the last decade, mass spectrometry (MS)-based immunopeptidomics has emerged as a powerful analytical tool for comprehensive profiling of the immunopeptidome (2, 7), enabling the discovery of bacterial antigens presented during infection (8–16) or tumor-specific epitopes that can be used for immunotherapy (17–21), among other applications.

A typical MS-based immunopeptidomics workflow consists of three steps. It starts with sample preparation, which comprises the pull-down of MHC-peptide complexes and elution of purified immunopeptides. This is followed by liquid chromatography-tandem mass spectrometry (LC-MS/MS) analysis, where mass spectra of the intact and fragmented immunopeptides are recorded. Lastly, MS data is analyzed by search engines and software tools to match the acquired spectral data to corresponding peptide sequences (22–25). While all three steps influence the quality of the final results, robust sample preparation is key as it directly influences the net peptide yield and overall reproducibility. This step is the most time-consuming due to the complexity of immunopeptide isolation and is often considered a major bottleneck for the clinical implementation of immunopeptidomics pipelines (26). Sample preparation begins with mild lysis of cells or tissues to extract MHC-peptide complexes in their native state. Immunoprecipitation (IP) is the preferred method to isolate these complexes, using specific antibodies for MHC-I and MHC-II. Once isolated, immunopeptides are eluted from the MHC molecules under mild acidic conditions, followed by a desalting step to purify the peptides for LC-MS/MS analysis (22, 27, 28). An alternative, antibody-free approach for MHC-I peptide isolation is mild acid elution (MAE) directly from the cell surface (29–31).

In recent years, technological advances have revolutionized the field of immunopeptidomics, allowing high-throughput extraction and analysis of immunopeptides. Innovations in MS instrumentation and data acquisition have greatly improved the sensitivity of immunopeptidomics workflows. This includes for instance trapped ion mobility spectrometry (TIMS) (28, 32, 33), data-independent acquisition (DIA) (31, 34–36), and advanced data analysis by incorporating multiple search engines and/or data-driven rescoring (32, 33, 37, 38). Consequently, this has facilitated the evolution of sample preparation to higher throughput workflows on lower input samples. Initial sample preparation workflows, despite showing excellent peptide enrichment, required hundreds of millions of cells as starting material and depended on manual IP of immunopeptides in tubes or columns (15, 22, 25, 39). These workflows were laborious, time-consuming, and prone to variability, limiting their utility for large sample sets or scarce primary specimens. Since then, IP procedures have significantly improved and upgraded from individual column with large sample amounts to multi-well plate format compatible with smaller input (26, 40–42). Together with increased instrumental and computational sensitivity, such miniaturized sample preparation workflows have overcome the challenge of performing immunopeptidomics on samples with limited material such as scarce tumor sections, lymph node biopsies, and rare cell populations.

At the same time, automation in sample preparation has remarkably enhanced overall throughput, reproducibility and sensitivity by reducing sample loss and contamination risks (26, 28, 40–46). The use of 96-well plates in immunopeptidomics sample preparation was first introduced by Chong *et al.* (26) in 2018, using stacked 96-well filter plates in combination with a positive pressure device for sequential MHC-I and MHC-II IP on tissue and cell lysates (15, 25). Since then, several other advancements in immunopeptidomics sample preparation have been reported (19, 27, 29–32), focusing on further optimizing throughput in 96-well plate workflows (28, 44) or maximizing sensitivity for low-input samples, for instance, using microfluidics (42, 43, 45, 46). However, most workflows today focus on either throughput or sensitivity, while ideally, the combination of both should be achieved. To fill this gap, we have developed a refined sample preparation workflow that is fully optimized to process low-input and crude samples in 96-well plates, aiming to combine high throughput with maximal sensitivity. We implemented a miniaturized and automated workflow in a 96-well format using a Tecan Resolvex® A200 liquid handling positive pressure device and demonstrate sensitive detection of immunopeptidomics from 32 million to 0.5 million human cells, with the ability to recover nearly 300 predicted MHC-I binders from just 20,000 JY cells. We showcase how this miniaturized platform facilitates bacterial antigen discovery for *Listeria monocytogenes* and the tuberculosis vaccine strain Bacillus Calmette-Guérin (BCG) in a quantitative manner.

## EXPERIMENTAL PROCEDURES

### Cell lines

The Epstein–Barr virus (EBV)-immortalized human B cell line JY (ECACC 94022533), human HeLa cells (ECACC 93021013), and U937 (ATCC) cells were cultured at 37 °C in a humidified atmosphere at 10% CO_2_. JY and U937 cells were maintained in RPMI 1640 and GlutaMAX medium (#61870036, Thermo Fisher Scientific) with 10% heat-inactivated fetal bovine serum (FBS, #10270106, Thermo) without antibiotics. U937 cells were differentiated using phorbol 12-myristate 13-acetate (PMA) at 200 nM final concentration for 48 h. HeLa cells were grown without antibiotics in DMEM medium (#21969035, Thermo) supplemented with 10% FBS, 2□mM GlutaMax, 1% non-essential amino acids (#11140035, Thermo), 1□mM sodium pyruvate (#11360039, Thermo) and 10□mM HEPES (#15630056, Thermo). All the cells were tested negative for mycoplasma contamination. Cells were counted and volumes corresponding to the number of cells in the serial dilution experiment were collected by centrifugation at 300 ×□*g*, washed twice with ice-cold PBS and stored at -80 °C.

### Bacterial culture

A defrosted aliquot of BCG Pasteur strain was grown in Difco^TM^ Middlebrook 7H9 broth Medium (Becton Dickinson) supplemented with 10% oleic albumin dextrose catalase (OADC), 0.2% glycerol, and 0.05% Tween^®^80 (Sigma-Aldrich, Merck), shaking at 37 °C. *Listeria monocytogenes* EGD (BUG600 strain) was grown in brain heart infusion (BHI) broth (#10462498, Thermo), shaking at 37 °C. All the BCG and *Listeria* work was carried out aseptically in a biosafety level 2 facility.

### BCG and *Listeria* infection of U937 cells

U937 cells were differentiated into phagocytic macrophage-like cells by stimulation with 200 nM PMA for 48 h, followed by 24 h of rest in PMA-free media. BCG was cultured into the log phase and pelleted by centrifugation at 3900 x *g* for 7 min. Bacteria were washed once with PBS, resuspended in PBS, and filtered through 25, 26, and 27 gauge needles to acquire a single-cell suspension. Differentiated U937 cells were infected at a multiplicity of infection (MOI) of 20 for 3 h, and the medium was aspirated to remove extracellular BCG and after which fresh media was added. Infected and uninfected cells were harvested with a cell scraper after 24 h, washed two times with PBS, and dry cell pellets were stored at -80 °C.

*Listeria* was grown overnight and diluted in BHI to a density of 1e9 bacteria/mL, washed twice with PBS and resuspended in U937 growth medium without FBS. U937 cells were washed two times with PBS prior to 1 h infection with *Listeria* at an MOI of 25. Following infection, the cells were washed twice with PBS and further cultured for 23 h in cell culture medium supplemented with 10% FBS and 40□µg/mL gentamicin (#G1397, Sigma-Aldrich, Merck) to eliminate extracellular bacteria. Cells were harvested using a cell scraper, washed two times with PBS, and dry cell pellets were stored at 80 °C.

### Generation of immunoaffinity columns for MHC Class I and II pull-down

W6/32 monoclonal pan-MHC class I and PdV5.2 monoclonal pan-MHC class II antibodies were purified from HB95 (HB-95™, ATCC) and PdV5.2 (kindly provided by Bart Vandekerckhove, Ghent University (47)) hybridoma cells, respectively. To prepare the W6/32 immunoaffinity column, 3 mg of antibody was diluted in 5 mL tris-buffered saline (TBS) and added to 1 mL pre-washed protein-A Sepharose 4B packed beads (#101041, Thermo) in glass Econo-columns® (#7374150, Bio-Rad). To prepare the PdV5.2 immunity column, 1.5 mg of antibody was diluted in 5 mL TBS and added to protein-G Sepharose 4B packed beads (#101242, Invitrogen). Antibodies were incubated at room temperature for 1 h on an Ika® rolling tube mixer device. Each column was washed with 0.2 M sodium tetraborate buffer (pH 9, #B3545, Sigma-Aldrich, Merck) followed by 40 min chemical crosslinking using 20□mM dimethylpimelimidate (#D8388, Sigma-Aldrich, Merck), freshly dissolved in sodium tetraborate buffer. After crosslinking, antibody-bead complexes were washed and incubated for 2 h with 0.2 M ethanol amine (pH 8, #149582500, Thermo) to quench the crosslinking reaction. A more detailed description of the procedure can be found in the Supplementary Information.

### High-throughput isolation and purification of MHC-I and MHC-II peptides

#### Cell lysis

Cell pellets ranging from 0.5-32 million cells were lysed in 0.1 mL mild lysis buffer containing 1% octyl-β, D-glucopyranoside (OGP) (#O9882, Sigma-Aldrich, Merck), 0.25% sodium deoxycholate (#1065040250, Millipore, Merck), 1.25× cOmplete protease inhibitor cocktail (#4693159001, Roche), 1□mM phenylmethylsulfonyl fluoride (PMSF) (#52332, Sigma-Aldrich, Merck), 0.2□mM iodoacetamide (IAA) (#I1149, Sigma-Aldrich, Merck), and 1□mM ethylenediaminetetraacetic acid (EDTA) (#EDS, Sigma-Aldrich, Merck) in 150 mM NaCl, 50 mM Tris (pH 8). Cells were lysed for 1 h on ice while resuspending every 15 minutes, then centrifuged at 20,000 × *g* at 4 °C for 10 min to remove cell debris, followed by a second centrifugation at 20,000 × *g* at 4 °C for 30 min to further clear the lysate.

We utilized a Resolvex A200 positive pressure processor (Tecan) to automate the washing and elution steps. Subsequent immunoprecipitations (IPs) of MHC-I and MHC-II complexes were performed using a single-use 96-well filter microplate (0.7 µm glass fiber membrane, 2 mL/well, long drip Agilent [part #201719-100]). Approximately 5-10% low positive pressure was applied at each step to maintain a uniform liquid flow through the plates. A more detailed description of the procedure can be found in the Supplementary Information.

#### Preparation of filter plates for MHC-I and II pulldown

Before IP, plates were activated with 1 mL 100% acetonitrile (ACN) (Sigma-Aldrich) twice, followed by three times 1 mL 0.1% trifluoroacetic acid (TFA) and finally with three times 1 mL 150 mM NaCl in 50 mM Tris (pH 8). Next, the MHC-I and MHC-II crosslinked beads were loaded on their respective plates (further referred to as “MHC-I filter” and “MHC-II filter” plates, Fig. 1A). Beads were washed five times with 1 mL 150 mM NaCl in 50 mM Tris (pH 8).

**Figure 1.**
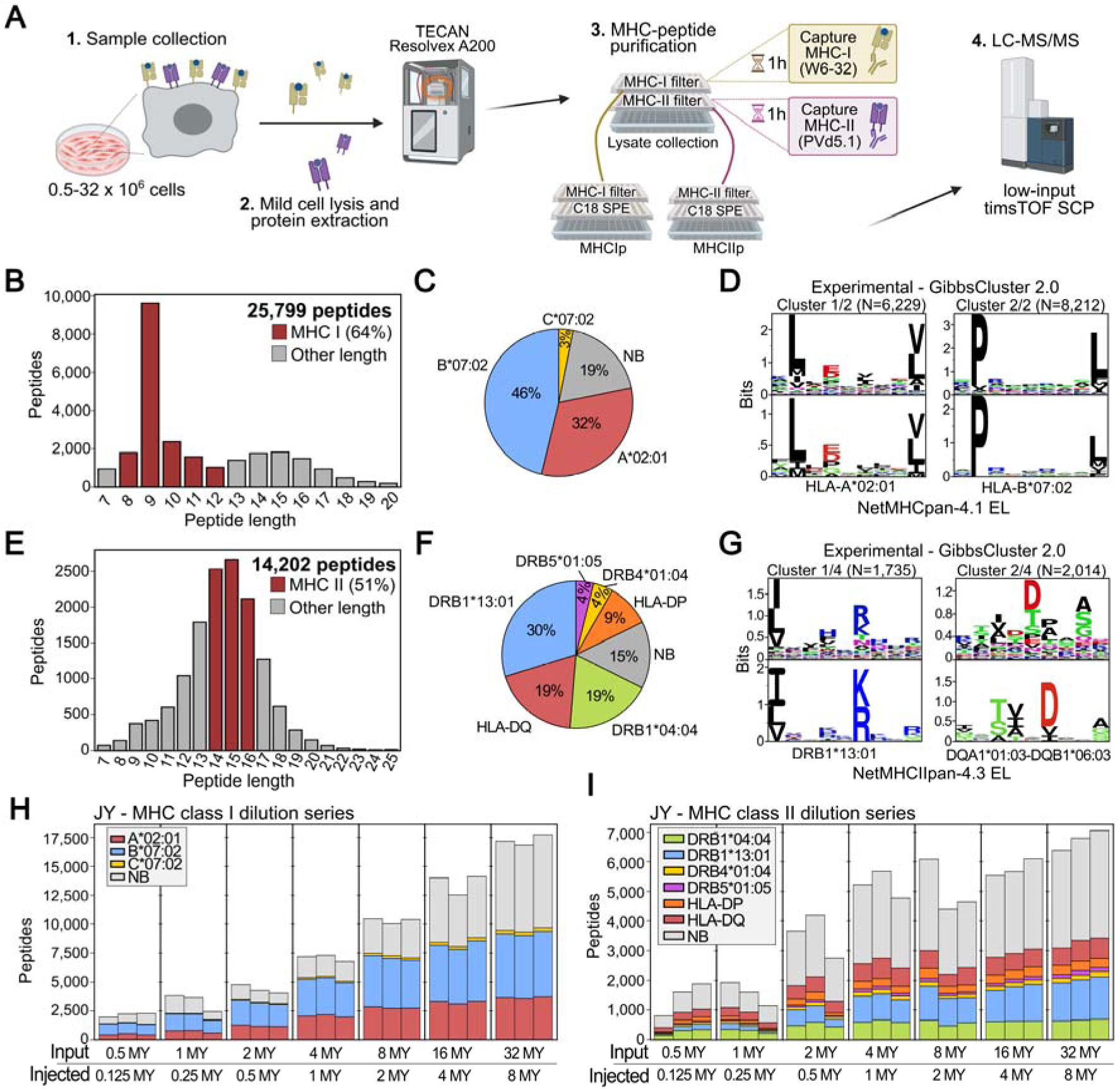
A sensitive automated workflow for high throughput immunopeptidomics. (**A**) General overview of the workflow. Cells are lysed in mild conditions, after which proteins are extracted and loaded on 96-well plates with pre-washed anti-MHC immunoaffinity beads. Sequential MHC-I and MHC-II IP, elution and desalting are performed in a semi-automated fashion on a Resolvex A200 (Tecan), after which peptides are collected and 25% is injected for LC-MS/MS on a timsTOF SCP instrument. Figure created with BioRender.com. (**B**) Histogram showing the length distribution of peptides identified after MHC class I pull down. Peptides compatible with MHC-I binding (length 8-12) are indicated in red. (**C**) Proportion of identified 8-12 mers (peptide Q-value < 1%) predicted to bind to the indicated HLA alleles by NetMHCpan-4.1 (53). (**D**) Unsupervised Gibbs clustering (55) of identified 8-12 mers resulting in sequence logos that match MHC class I eluted ligand (EL) motifs of NetMHCpan-4.1 (53). (**E**) Histogram showing the length distribution of peptides identified after MHC class II pull-down and elution. Peptides compatible with MHC-II binding (length 14-16) are indicated in red. (**F**) Proportions of identified 14-16 mers predicted to bind to the indicated HLA alleles by NetMHCIIpan-4.3 (54). For HLA-DQ and HLA-DP, the possible α–β chain pairings (*e.g.* HLA-DQA10103-DQB10603) were grouped together. (**G**) Unsupervised Gibbs clustering (55) of identified 14-16 mers resulted in sequence logos that match MHC class II eluted ligand (EL) motifs of NetMHCIIpan-4.3 (54). (**H-I**) Number of identified peptide sequences per replicate sample grouped by cell input amount. Samples from each amount were processed and searched together, and 25% of the peptide material was injected on a timsTOF SCP. Peptides are colored according to binding prediction for JY MHC class I alleles (**H**) and class II alleles (**I**).

#### Subsequent MHC-I and MHC-II immunoprecipitation

Lysates were added to the MHC-I filter plate and incubated for 1 h at 4 °C to capture MHC-I complexes. Next, the unbound lysate fraction of the MHC-I IP was collected using the Tecan and loaded on the MHC-II filter plate, followed by 1 h incubation at 4 °C. Meanwhile, the MHC-I filter plate was washed five times with 1 mL 150 mM NaCl in 50 mM Tris (pH 8) using the Tecan. After 1 h incubation, the MHC-II filter plate was washed similarly.

#### Immunopeptide desalting and elution

Sep-Pak tC18 plates (#186002321, Waters) were activated with two times 1 mL 100% ACN and three times 1 mL Milli-Q water containing 0.1% TFA. Each MHC filter plate was stacked on top of a Sep-Pak plate (Fig. 1A) and MHC-peptide complexes were eluted with 0.2 mL 10% acetic acid five times. Next, we removed the filter plate and washed the Sep-Pak plate three times with 1 mL of 0.1% TFA. Thereafter, a collection plate (#201240-100, Agilent) was mounted under the Sep-Pak tC18 plate. MHC-I peptides were eluted three times with 25% ACN, while MHC-II peptides were eluted with 40% ACN. The eluted immunopeptides were then transferred to protein LoBind tubes (#0030108450, Eppendorf), vacuum dried and stored at 20°C until LC-MS/MS injection.

### LC-MS/MS analysis

Dried peptides were dissolved in 50 μL 0.1% ACN/0.1% TFA from which 12.5 μL was injected for LC-MS/MS analysis on a 3000 RSLC nanoLC in-line connected to a timsTOF SCP mass spectrometer (Bruker). Of note, the dried peptides from low input dilution series (Fig.2) were dissolved in 17 μL 0.1% ACN/0.1% TFA from which 15 μL was injected for LC-MS/MS analysis. Trapping was conducted at a flow rate of 20 μL/min for 2 min in loading solvent A on a 5 mm trapping column (Thermo, Pepmap, 300 μm internal diameter [I.D.], 5 μm beads). The sample was then separated on a reverse-phase column (Aurora elite 75µm x 150 mm 1.7 μm particles, IonOpticks) following elution from the trapping column. Separation was achieved using a linear gradient, starting at 0.5% MS solvent B (0.1% FA in water/ACN 20:80, v/v) at 250 nL/min for 30 min, increasing to 37.5% MS solvent B, then reaching 55% at 38 min, and further rising to 70% at 40 min. This was followed by the wash for 5 min and re-equilibration with 99.5% MS solvent A (0.1% FA in water). The flow rate was adjusted from 250 nL/min to 100 nL/min at 20 minutes and restored to 250 nL/min at 40 min. A ten PASEF/MSMS scan acquisition method was employed in DDA-PASEF mode with a precursor signals intensity threshold of 500 arbitrary units. Precursors were isolated with a 2 Th window below *m/z* 700 and 3 Th above, with active exclusion for 0.4 min upon reaching a target intensity threshold of 20,000 arbitrary units. In case of MHC-I peptides, an adapted “Thunder” polygon (32) in the *m/z*-IM plane (range 0.7 to 1.25 V s/cm^2^) was applied to include HLA-I singly charged precursors, while a broad polygon (range 0.75 to 1.65 V s/cm^2^) was used for MHC-II peptides (**Supplemental Fig. S1**). The mass spectrometer was operated in non-sensitive mode with an accumulation and ramp time of 100 ms, analyzing in MS from 100 to 1,700 *m/*z, applying collision energy according to the IM range specified in **Supplemental Table 1**.

**Figure 2.**
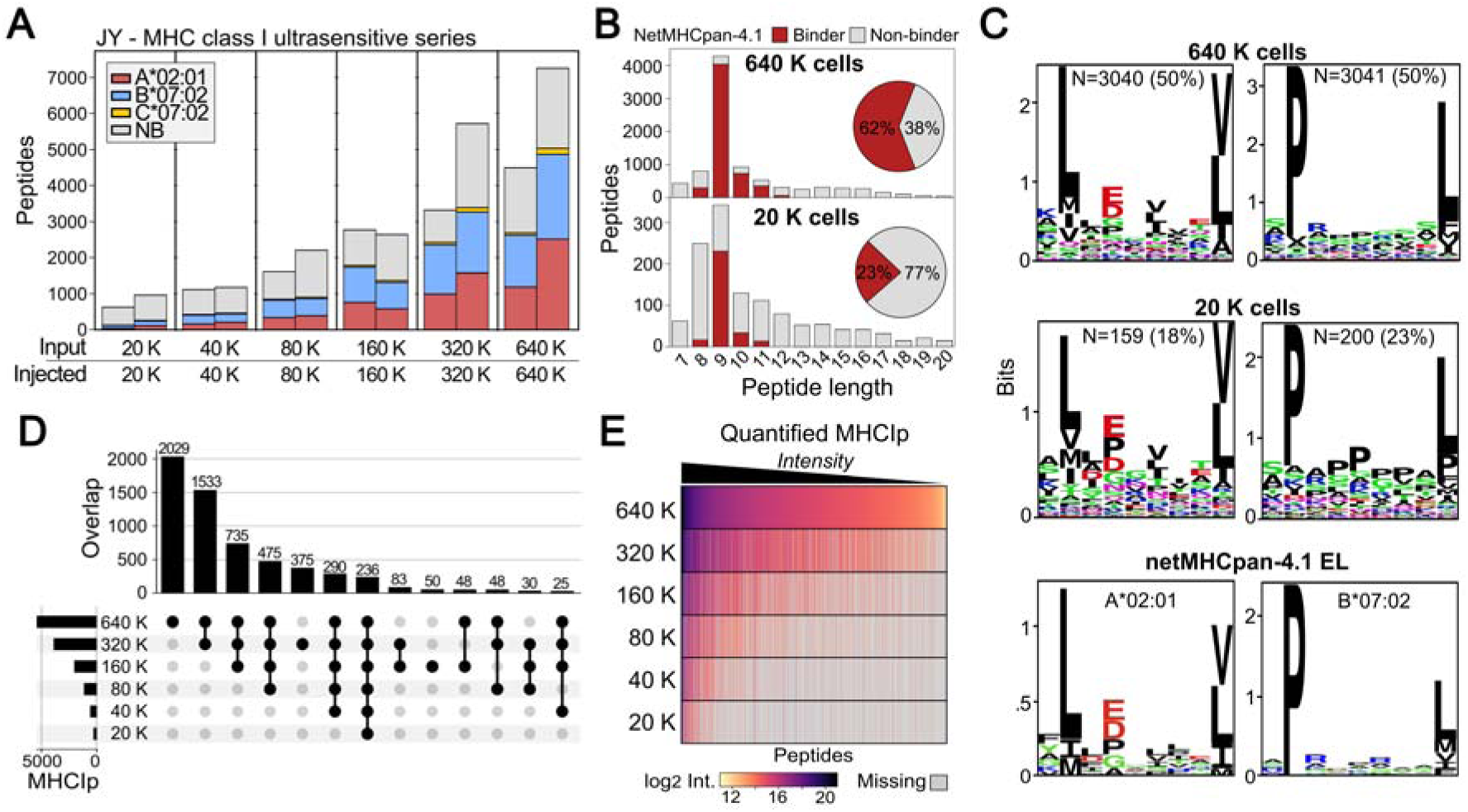
Towards ultrasensitive immunopeptidomics. (**A**) Number of identified peptide sequences per sample grouped per cell dilution series from 640,000 (640 K) down to 20,000 cells (20K). Samples from each input amount were processed and searched independently, and 100% of the isolated immunopeptides were analyzed by LC/MS-MS. Peptides are colored according to MHC binding prediction by NetMHCpan-4.1 (53). (**B**) Histogram showing the length distribution of peptides identified after MHC class I pulldown and elution from 640 K and 20 K cells. Peptides compatible with MHC-I binding (length 8-12) are indicated in red. (**C**) Unsupervised Gibbs clustering (55) of identified class I peptides from 640 K and 20 K cells (*top and middle*) resulted in sequence logos that match NetMHCpan-4.1 eluted ligand (EL) sequence logos for A*02:01 and B*07:02 (*bottom*). (**D**) UpSet plot showing the overlap of identified predicted MHC-I peptides (MHCIp, NetMHCpan-4.1 %Rank < 2) across the different input amounts. Only intersections of more than 10 peptides are displayed. (**E**) Heatmap showing log_2_ intensities of all peptides identified in the 640 K dilution, sorting peptides from highest to lowest intensity in this dilution (*right to left*). Missing values are colored in grey. Dilutions were searched separately using the FragPipe ‘Nonspecific-HLA’ workflow.

### Data analysis and processing

#### Immunopeptidomics searches and analysis

All raw proteomics data were searched using a multi-search engine workflow with follow-up rescoring as described in Willems *et al.* (33). In brief, timsTOF (.d) data was searched by four search engines (MSFragger (48), Comet (49), Sage (50) and PEAKS Studio 12 (51)) after which search results for each engine were rescored in parallel using TIMS^2^Rescore (52). Peptides identified below a 1% peptide q-value were reconciled at the spectrum-level, excluding the rare cases where a spectrum was matched to distinct peptides by different search engines (33). Downstream immuno-informatic analyses included NetMHCpan-4.1 (53) and NetMHCIIpan-4.3 (54) prediction of binding strength for 8–12-mers and 14–16-mers, respectively, and unsupervised Gibbs clustering (55). Default NetMHCpan %Rank score thresholds were used to define strong and weak MHC class I binders (below 0.5 and 2, respectively) and MHC class II binders (below 2 and 5, respectively).

#### Differential analysis

To analyze differential immunopeptide abundances between infected and uninfected samples, we used FragPipe (version 22.0) including MSBooster (38) and IonQuant quantification (56). Resulting peptide intensities (from ‘combined peptides.tsv’ reports) were used for peptide-level differential statistics. To assess differential peptide abundance, we resorted to an empirical Bayes test in limma (57) following recommendations of a statistical benchmark (58). Significantly regulated peptides were filtered at an adjusted *P* value ≤ 0.05 and absolute fold change ≥ 2. This differential test does not impute missing values, with the drawback of not reporting peptides exclusively detected in one of the conditions (infected/uninfected). Therefore, we performed an additional differential detection analysis where we discerned peptides/proteins uniquely identified in one condition (three out of four replicates) and absent in the other condition.

#### Gene set enrichment analysis

For the differential immunopeptides identified during infection, their corresponding UniProtKB accessions were used for protein-level gene set enrichment analysis. In the case of a peptide matched multiple proteins, the first UniProtKB accession was used as representative. Accessions were entered in the online webtool g:Profiler using default settings (59).

#### Visualisation

Plots were generated using matplotlib (60) and seaborn (61) in Python (version 3.8.10). Sequence logos were generated using Logomaker (62). UpSet plots were made using the Python implementation module *upsetplot* (63).

## RESULTS

### An automated workflow for high-throughput immunopeptidomics

Our previous immunopeptidomics workflow required approximately 500 million cells as input for manual immunoprecipitation (IP) in large columns (8, 15), thus requiring large sample input amounts with limited throughput. Here, we aimed to miniaturize and automate sample preparation by reducing the input amount 10- to 50-fold while increasing sample throughput by using a 96-well format. We initially tested optimal conditions for cell lysis, IP and elution on JY cells, an EBV-immortalized human B cell line, and found that static IP at high protein concentration (small lysate volume) with elution in 10% acetic acid led to the highest number of identified MHC-I and II peptides (**Supplemental Fig. S2-S5**). This optimized procedure was then automated on a Tecan Resolvex® A200 positive pressure device, ensuring a controlled and consistent flow through the wells. In the resulting workflow, cell pellets are lysed in 100 µL lysis buffer and lysates are loaded on 96-well filter plates for sequential pull-down of MHC-I and II complexes, followed by acetic acid elution and Sep-Pak C18 purification of immunopeptides, all in an automated fashion (**Fig. 1A**). Performing MHC-II after MHC-I pulldown was necessary to avoid co-enrichment of MHC-I peptide complexes during MHC-II IP (**Supplementary Fig. S6**). We also found that the pan-MHC-II PdV5.2 antibody shows similar performance to other MHC class II antibodies (TU39 or IVA12, **Supplemental Fig. S6**). Finally, isolated immunopeptides are analyzed by LC-MS/MS analysis using a timsTOF SCP mass spectrometer operating in DDA-PASEF mode, followed by immunopeptide identification using four different search engines with data-driven rescoring as described (33).

To test the sensitivity of the immunopeptidomics platform, we enriched MHC class I and II peptides from JY cells, using input amounts ranging from 32 to 0.5 million cells. For each input amount, triplicate samples were processed side by side, and from each sample, 25% of the isolated immunopeptides were injected onto the timsTOF SCP. First, we assessed the global quality of the immunopeptidome by searching all samples together. Over all input amounts, a total of 25,799 peptides were identified in the MHC-I pulldown, including 16,404 (64%) 8- to 12-mers (**Fig. 1B, Supplemental Data S1**). The majority of these 8- to 12-mers (81%) were predicted to bind at least one JY HLA allele (**Fig. 1C**) and clustered into matching peptide sequence motifs (**Fig. 1D**). For MHC-II, we identified 14,202 peptides in JY cells, including 7,304 (51%) 14- to 16-mers (**Fig. 1E, Supplemental Data S2**), of which 85% were predicted binders (**Fig. 1F)** and that yielded the expected MHC sequence motifs (**Fig. 1G**).

Next, we searched the samples from each input amount separately for fair comparison. For both MHC class I and II pulldowns, we observed a progressive decline in the number of identified peptides with decreasing input amount (**Fig. 1H-I**). For instance, from 0.5 million cells on average 2,158 MHC-I peptides were identified, while from 32 million cells on average 17,266 peptides were identified, approximately eight times more. Interestingly, the proportion of NetMHCpan-predicted MHC-I binders is slightly higher at lower input numbers. With 0.5 million cells as input, 66% of the peptides were predicted to bind JY HLA alleles on average, while for 32 million cells, 55% were predicted binders. As such, an increase in cell number input is not linearly correlated with the number of identified MHC class I-binding peptides. This is exemplified by the fact that the number of predicted MHC class I binders starts to plateau around 16 million cells (**Fig. 1H**), a trend that was also observed for MHC class II peptides (**Fig. 1J**). We also observed that for both MHC-I and II peptides, the median binding strength increased with decreasing input amounts, indicating that the identification of strong-binding peptides is favored over weak-binding peptides in low-input samples (**Supplemental Fig. S7**).

To test whether these observations generalize beyond JY cells, we also isolated MHC-I peptides from different amounts of HeLa and U937 cells, representing human epithelial and macrophage cells, respectively. The detected peptides were also of high quality and showed preferred binding patterns to the anticipated HLA alleles, with a higher proportion of 8-mers in U937 cells due to immunopeptide presentation on HLA-B*18:01 and HLA-B*51:01 (**Supplemental Fig. S8A-F, Supplemental Data S3-4**) (64). While in general fewer peptides were identified compared to JY cells, for both cell types the number of identified immunopeptides also plateaued with a sample size of 16 million cells, which we therefore defined as the optimal input amount for maximum detection of immunopeptides (**Fig. 1 H-I**, **Supplemental Fig. S8H-I**). Together, these experiments established an automated workflow for immunopeptidomics sample preparation in 96-well format, compatible with a wide range of input amounts.

### Towards ultrasensitive immunopeptidomics

Given the successful identification of more than 2,000 MHC class I peptides from 0.5 million cells (**Fig. 1H**), we tested the capability of our platform to identify immunopeptides from very low (< 1 million) cell input amounts. Starting from 0.64 million cells, we generated a two-fold serial dilution series down to 20,000 cells as input. The same automated sample preparation workflow was used, but now the full sample amount (100%) was injected on the timsTOF SCP. Analyzing both samples with 640,000 cells, we identified 8,858 unique peptides of which 5,482 (62%) were predicted MHC class I binders (**Fig. 2A-B, Supplemental Data S5**). At a 30-fold lower input of only 20,000 cells, we still detected 1,244 peptides with 292 (23%) predicted binders. A peak of 9-mers was still discernible, and unsupervised Gibbs clustering recovered the leading JY HLA sequence motifs of HLA-A*02:01 and HLA-B*07:02 (**Fig. 2B-C, Supplemental Fig. S9**), indicating successful immunopeptide isolation at these ultra-low inputs. In addition, we repeated this experiment but now with application of the MS-compatible nonionic surfactant n-dodecyl-β-D-maltoside (DDM), which is used to increase peptide identification in single-cell experiments by decreasing peptide adsorption to plasticware (65). The same trend and number of identified peptides was observed without DDM, suggesting no beneficial effect of DDM in this setup (**Supplemental Fig. S10, Supplemental Data S6**).

Comparing identified MHC binding peptides across input amounts revealed a common ‘core’ subset of class I peptides that progressively narrows with each dilution, eventually reaching 236 peptides that were identified across all dilutions (**Fig. 2D**). Thus, 236 out of 292 (81%) MHC-I-presented peptides in the 20,000-cell samples were present in all dilutions. To address why these particular immunopeptides were retained at very low input, we explored multiple biological and technical reasons. To explore the role of peptide intensity, we ran a FragPipe HLA peptidome analysis with IonQuant (56) quantification separately for each dilution (**Supplemental Data S7**). This revealed MS intensity to be a determining factor, with high-abundant immunopeptides in the 640,000 cell input more likely to be identified in lower dilutions (**Fig. 2E**). The binding strength of predicted MHC-I binders was similarly distributed across dilutions (**Supplemental Fig. S11**). We also inspected the potential sources of the relatively higher proportion of predicted non-binding peptides at very low cell dilutions. While the origin of most of these peptides remains unclear, we see that 25 to 30% of these peptides belong to typical proteomic contaminants (e.g. keratins) (**Supplemental Fig. S12**).

### Detection of bacterial immunopeptides presented by infected cells

When applied to infected cells or tissues, immunopeptidomics allows the untargeted detection of bacterial immunopeptides, making it a powerful technology for antigen discovery and bacterial vaccine development (8, 66). We previously reported the detection of bacterial immunopeptides presented by cells infected with *Listeria monocytogenes* (8, 15, 33, 34) using several hundred million cells as input. Here, we aimed to test whether our optimized workflow would achieve similar performance in bacterial antigen detection using lower cell input amounts. To this end, we infected 16 million U937 cell samples with *Listeria monocytogenes* EGD (further referred to as *Listeria*) and isolated MHC class I immunopeptides from three infected and three uninfected samples for LC-MS/MS analysis (**Fig. 3A**). In total, 20,008 peptides were identified, including 12,995 8- to 12-mers of which 7,922 (61%) were predicted binders of U937 HLA alleles (**Fig. 3B, Supplemental Data S8**). The detected peptides showed preferred binding patterns to the anticipated HLA alleles, with again a higher proportion of 8-mers resulting from B*18:01- and B*51:01-presented peptides (**Supplemental Fig. S13A**). Next, we stringently filtered high-confidence bacterial peptides as described (33). While most peptides were of human origin, we detected 50 high-confidence bacterial peptides derived from 35 *Listeria* proteins. Of these, elongation factor Tu (tuf), virulence factors hly/LLO and inlC, periplasmic oligopeptide-binding protein OppA and cell wall carboxypeptidase LMON_2776 were represented by multiple immunopeptides (**Fig. 3C**). This observation is in line with the previously reported immunodominance of these *Listeria* antigens, where tuf, hly/LLO and OppA were also identified by multiple immunopeptides in infected HeLa and HCT-116 cell cultures (8, 33, 34). In addition to *Listeria*, we applied our workflow to U937 cells infected with BCG. Here, 12,833 MHC class I peptides were identified, including 8,480 8- to 12-mers with 4,992 (59%) predicted binders (**Fig. 3D, Supplemental Fig. S13B, Supplemental Data S9**). We detected 41 high-confidence bacterial peptides derived from 32 BCG proteins. Again, several BCG proteins were represented by multiple immunopeptides, including elongation factor Tu (tuf), groEL2, groES and hupB (**Fig. 3E**). Interestingly, peptides from these three proteins were also identified by a previous immunopeptidomics study that mapped MHC class II peptides on BCG-infected cells (9, 10), and the same holds true for embC (67). Also IniB was identified in *Mycobacterium tuberculosis*-infected macrophages (68). Together, these data demonstrate the ability of our workflow to effectively capture relevant bacterial immunopeptides from limited amounts of infected cells.

**Figure 3.**
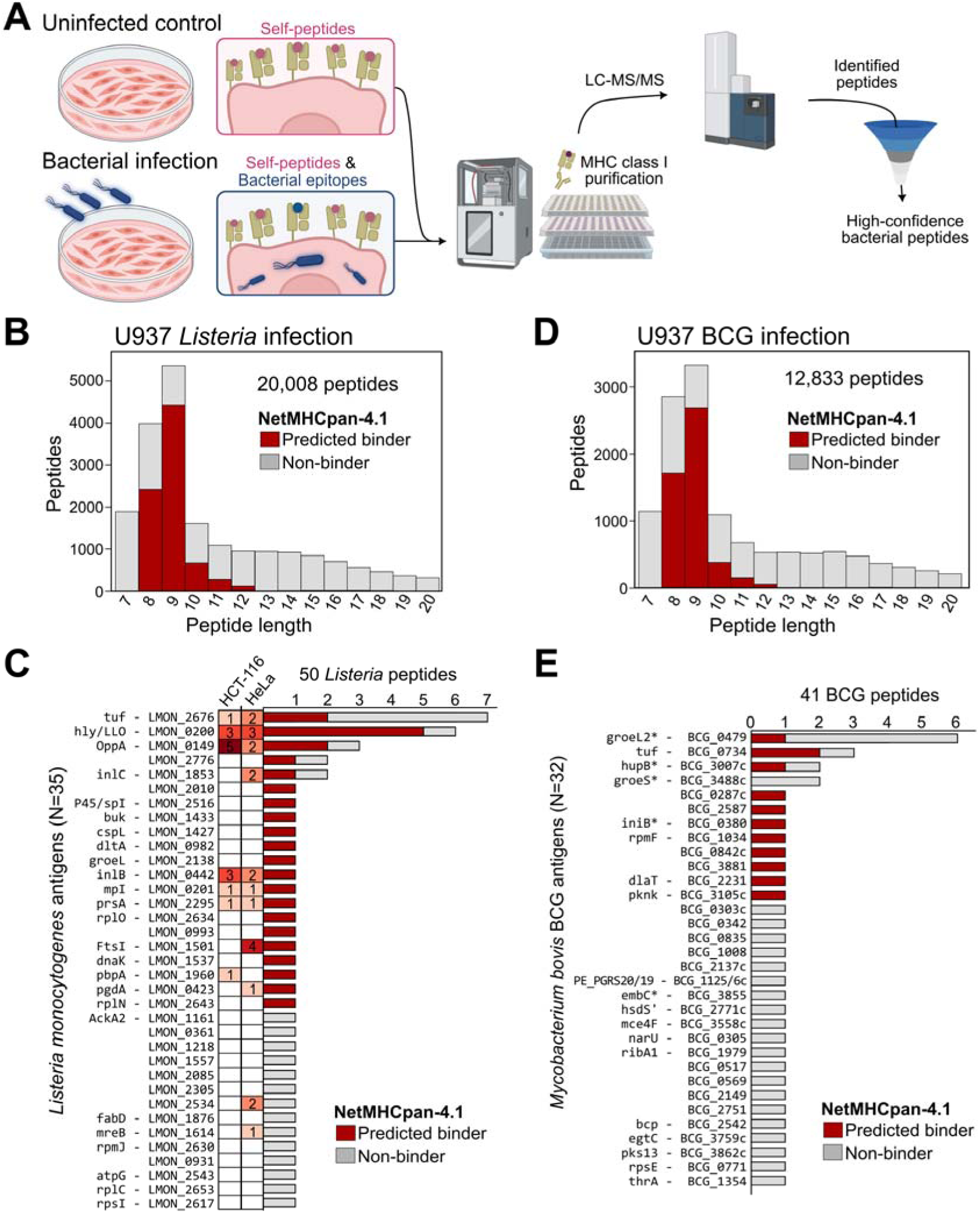
Bacterial antigen detection. (**A**) Bacterial immunopeptidomics workflow scheme. Differentiated U937 cells infected with bacteria and uninfected counterparts are subjected to the immunopeptidomics workflow. This enables the detection of bacterial epitopes among numerous host self-peptides. After purification and LC-MS/MS of immunopeptides, data was filtered to retain only high-confidence bacterial immunopeptides. Figure created with BioRender.com. (**B, D**) Histogram showing the length distribution of peptides identified after MHC class I pull down. Peptides predicted to bind to U937 MHC-I alleles by NetMHCpan-4.1 (53) (%Rank < 2) are indicated in red. (**C, E**) The number of unique immunopeptides identified per *Listeria* protein (**C**) or *Mycobacterium bovis* BCG protein (**E**) is shown in a histogram. Immunopeptides predicted as MHC class I binders by NetMHCpan-4.1 (53) (%Rank < 2) are indicated in green, other peptides in grey. (**C**) The number of identified peptides per cell line in our previous *Listeria* study (8) is displayed in a heatmap. (**E**) BCG proteins identified in previous immunopeptidomics studies (9, 10, 67, 68) are indicated by an asterisk (*).

### Quantitative remodeling of the human immunopeptidome upon BCG infection

Bacterial infection impacts many cellular processes, resulting in altered presentation of host cell immunopeptides in addition to bacterial peptides. Using a quantitative, nonspecific HLA workflow in FragPipe, we aimed to identify differential human immunopeptide presentation upon BCG infection. Firstly, we assessed the reproducibility of peptide quantification. Uninfected controls displayed a higher quantitative reproducibility (Pearson R = 0.885 ± 0.034) than BCG-infected samples (Pearson R = 0.794 ± 0.045), suggesting that infection introduces greater quantitative variability (**Supplemental Fig. S14A**). Further, the multidimensional scaling (MDS) plot based on immunopeptide intensities revealed a clear separation between uninfected controls and BCG-infected samples (**Supplemental Fig. S14B**). Next, we compared the intensities of the human immunopeptides between the BCG-infected and uninfected samples. This analysis revealed 1,301 immunopeptides that were significantly upregulated (adj. *P* ≤ 0.05, ≥ two-fold change) or uniquely present in the infected samples, while 117 peptides were upregulated or uniquely present in the uninfected samples (**Fig. 4A, Supplemental Data S10**). Gene set enrichment analysis revealed that many regulated peptides (both up and down) were derived from ribosomal proteins and many peptides upregulated upon infection originated from proteins linked to infection, inflammation and immune signaling (**Fig. 4B**). For instance, interleukin-1 beta (IL-1β) showed a drastic upregulation of 24 immunopeptides during infection, and was annotated alongside other immune-related proteins as part of the KEGG ‘IL-17 signaling pathway’ that was significantly enriched (adj. *P* 1.16×10^−4^) (**Fig. 4C**). IL-1β is a well-known pro-inflammatory cytokine that accumulates in monocytes and macrophages in response to pathogen infection (69, 70). Interestingly, it is produced in the cytosol as an inactive precursor protein that requires a maturation step for its release (70, 71), possibly facilitating its strong presentation by MHC-I under infection. In addition, proteins like Fcγ receptors (FCGR2A and B) and ARP2/3 complex subunits (ACTR and ARPC proteins) involved in ‘FcγR-mediated phagocytosis’ were more presented by infected cells (adj. *P* 0.041) (**Fig. 4D**). Taken together, these data show the high reproducibility and quantitative performance of our workflow, revealing immune remodeling under infection, with increased presentation by MHC class I molecules of proteins involved in inflammation, phagocytosis and other immune-related processes triggered by BCG infection.

**Figure 4.**
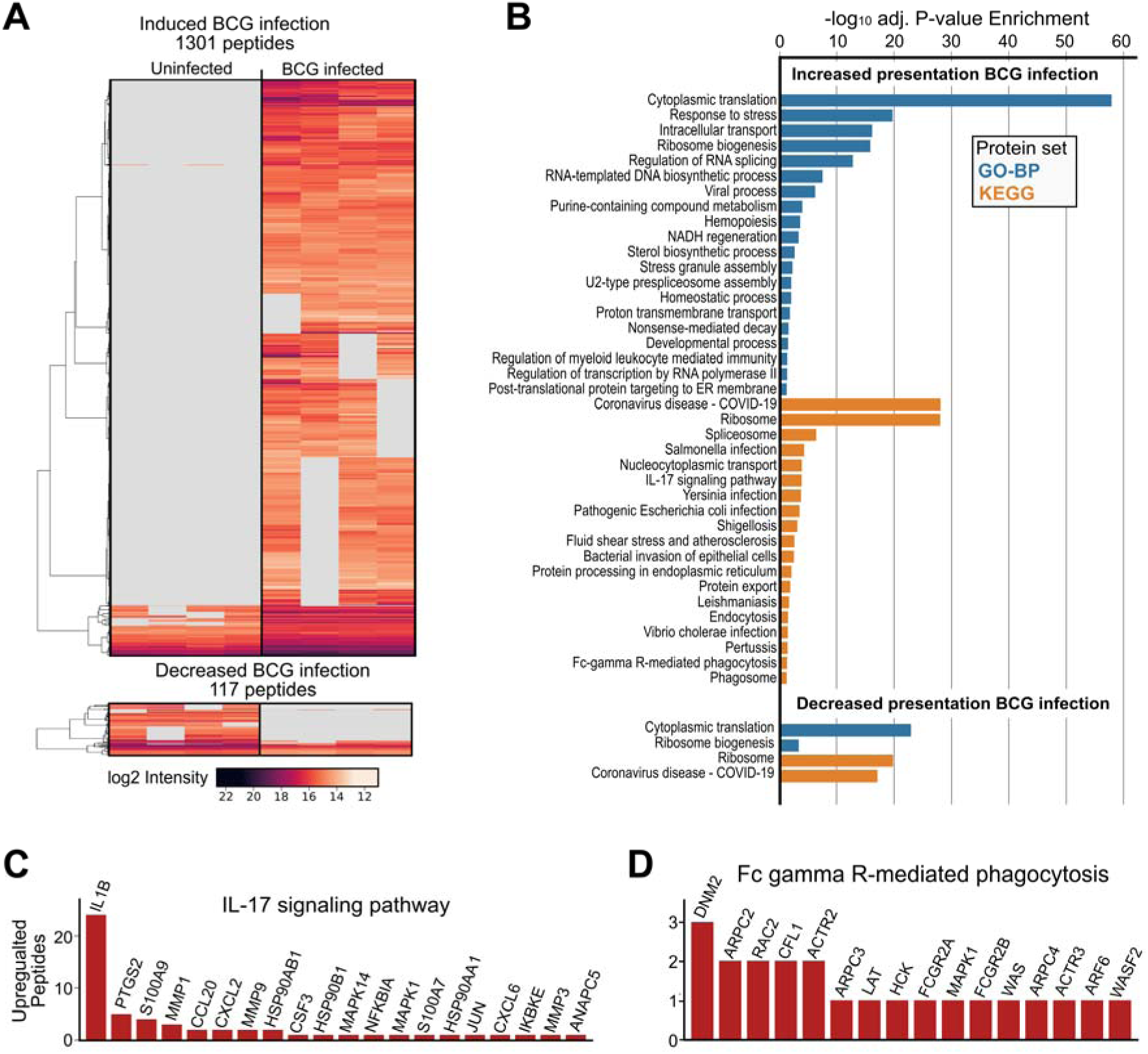
Host immunopeptide remodeling in response to BCG infection. **(A)** Heatmap showing log2 intensities of significantly differentially regulated immunopeptides (adj. *P* value ≤ 0.05 and absolute fold change ≥ 2) as well as immunopeptides uniquely detected in three out four replicates in infected or uninfected condition. (**B**) Functional enrichment analysis of UniProtKB protein accessions with up- or downregulated immunopeptides under infection using g:Profiler (59). (**C-D**) Number of significantly upregulated peptides in the enriched protein sets ‘IL-17 signaling pathway’ (**C**) and FcγR-mediated phagocytosis (**D**).

## DISCUSSION

We here describe a platform for immunopeptidomics sample preparation that is both sensitive and compatible with high-throughput analysis. Sequential capture of MHC class I and II immunopeptides is achieved through optimized IP in 96-well format on a positive pressure device. In combination with sensitive timsTOF SCP data acquisition and analysis (28, 32, 33) we demonstrate that the platform can comprehensively capture and identify MHC class I and II immunopeptidomes using sample sizes ranging from 32 to 0.5 million cells, with 16 million cells as optimum after which no further gain in detected binders was observed. Exploring the sensitivity limits of our platform, we could detect over 1,200 peptides from only 20,000 cells, albeit challenged with lower percentages of predicted binders. To further showcase the discovery potential of the platform, we performed bacterial antigen discovery in 16 million U937 macrophages infected with *Listeria* or BCG. This led to the identification of known and novel bacterial antigens, despite a 30-fold reduction in input material compared to previous analyses (8, 9, 33, 34, 67, 68). Thus, our optimized platform allows sensitive immunopeptide detection in a high-throughput fashion, enabling its application on biological sample types where input amounts are limited.

Cell lysis, IP and elution steps are determining steps for successful immunopeptide isolation, which were all carefully optimized. Traditional IP protocols typically mix antibody-bound beads with lysate under agitation (8, 25), however, we observed that MHC-I IP under static conditions produces equivalent results (**Supplemental Fig. S2A-C**). As a more important factor, we uncovered that efficient MHC-I IP heavily depends on a high protein concentration, which can be achieved by reducing the lysis buffer volume (**Supplemental Fig. S2D-E**). To increase sample throughput, we tested multiple 96-well filter plates with pore sizes ranging from 25 to 0.2 µm using a positive pressure device. With larger pores causing leakage during incubation steps and smaller pores requiring higher pressure, we selected a 0.7 µm filter plate as an optimal middle ground, allowing a steady flow without leakage nor clogging. Lastly, we tested IP elution and peptide purification methods (**Supplemental Fig. S4-5**), determining 10% acetic acid IP elution with Sep-Pak tC18 peptide purification as a preferred method.

Together, these optimizations led to the most important difference with previous multi-well sample preparation procedures (**Supplemental Table S2**): the small lysis volume used for IP of only 100 µl. Other multi-well procedures typically start from larger lysis volumes of 1 mL or higher such as the original 96-well procedure by Chong *et al.* (26). Inspired by this protocol, we also employ static IP, however, instead of relying on gravity flow, we optimized the plate pore size to prevent leakage and to allow flow control by positive pressure. Overall, our platform reduces sample transfers, thereby avoiding sample losses and enhancing sensitivity compared to procedures involving single tubes (8, 15) or incubation of deep well plates under rotation (28). Similar to other immunopeptidomics platforms using Waters (26), Agilent (40, 41) or KingFisher (42) devices, we implemented our procedure on a Tecan Resolvex A200 to automate the IP and immunopeptide elution. As an alternative approach, microfluidics set-ups have emerged that further miniaturize immunopeptidomics sample preparation with minimal sample transfers. However, these setups require highly specialized equipment and expertise for implementation while only processing a single sample each time, making this approach impractical for projects demanding higher-throughput.

For maximal performance, low-input sample preparation should be coupled to sensitive mass spectrometry data acquisition. To this end, we analyzed our samples on a timsTOF SCP instrument operating in DDA-PASEF mode, leveraging ion mobility-*m/z* peptide precursor selection polygons tailored for immunopeptidomics (28, 32, 33) (**Supplemental Fig. S1**). Importantly, the cell input numbers reported reflect the actual numbers used for lysis, more closely mimicking real-life applications compared to the cell equivalents that are sometimes used for method optimization (28, 32). However, it is important to note that immunopeptide identification numbers are largely dependent on the used cell line and their MHC expression levels, making comparisons across studies and cell types difficult. To further maximize reliable immunopeptide identification, we applied advanced data analyzed by integrating the results of four independent search engines that were rescored by TIMS^2^Rescore, (52) as recently described (33). In future work, we plan to couple our platform to alternative MS acquisition methods, including DIA-PASEF (29, 34, 72), or to other acquisition schemes such as Slice-PASEF (73) or Synchro-PASEF (74). Alternatively, the high sensitivity and speed offered by Orbitrap Astral instruments holds great promise for low-input immunopeptidomics (31). In terms of sample preparation, we envision to improve the sensitivity of our platform even further by using magnetic beads (42, 46) along with biotinylated MHC antibodies (36, 45) for IP, to work with even smaller sample volumes and to prevent endogenous antibody binding to protein-A or -G beads when processing tissue samples. Interestingly, mild acid elution (MAE) has re-gained interest and was shown to offer specific and sensitive enrichment of MHC class I peptides as a cost-effective alternative for antibody-based IP (29–31). However, due to its need for live cells in suspension, MAE has limited applicability to solid tissue samples and frozen cells (30, 75), though it was successfully applied to clinical tissue fragments (31). In addition, so far it has only been reported for MHC-I, limiting its applicability to samples where the immunopeptidome of both MHC classes are of interest.

In this study we have also demonstrated the ability of our immunopeptidomics platform to identify bacterial antigens in a lower-input setting using 16 million infected macrophages. This input downscaling substantially decreases the upfront cell culturing and hands-on time, which makes the procedure compatible with higher biosafety level pathogens for which the maximal number of cells that can be infected is limited. The lower input also allows quantitative immunopeptidomics analysis on multiple replicates and conditions. Firstly, we performed antigen discovery for *Listeria monocytogenes*, a foodborne model pathogen for which we previously gained a comprehensive view on its presented antigens, allowing us to cross-reference the *Listeria* antigens identified by the low-input platform. Despite a 30-fold reduction in input amount, we identified 50 MHC-I peptides from 35 *Listeria* proteins in infected U937 macrophages. Of these 35 antigens, 12 were identified in previous *Listeria* immunopeptidomics studies with high cell number inputs (8, 33, 34), including previously detected immunodominant antigens and virulence factors such as LLO, OppA, inlC and inlB. After this verification, we applied our platform on U937 cells infected with BCG, an attenuated *Mycobacterium bovis* strain. This analysis uncovered 41 MHC-I peptides from 32 BCG proteins, of which groEL2, groES, hupB and embC were also detected by previous BCG antigen discovery studies in higher input analyses (9, 10, 67), while iniB was picked up in *M. tuberculosis*-infected U937/A2 cells (68). Aside from the identification of pathogen-derived immunopeptides, bacterial infection can remodel the host immunopeptidome (14, 15). We previously reported how *Listeria* infection promoted presentation of immune-related self proteins in spleens of infected mice (15). Similarly, we here report how BCG infection of U937 macrophages leads to increased MHC-I presentation of host proteins partaking in immune pathways related to immune signaling and phagocytosis (**Figure 4**), pathways that were previously reported to be upregulated in BCG-infected macrophages (76). From a technical perspective, these analyses demonstrated the excellent quantitative reproducibility of our immunopeptidomics platform, with an average Pearson correlation of 0.88 between immunopeptide intensities in uninfected biological replicate samples. Together, these data validate our platform for biological immunopeptidomics applications where a combination of high sensitivity, high throughput and quantitative performance are required.

## Abbreviations

BCG: bacillus Calmette-Guérin
DDA: data-dependent acquisition
DIA: data-independent acquisition
EBV: Epstein-Barr virus
IP: immunoprecipitation
LC-MS/MS: liquid chromatography-tandem mass spectrometry
MHC: major histocompatibility complex
PASEF: parallel accumulation–serial fragmentation
TIMS: trapped ion mobility spectrometry.

## Data availability

The mass spectrometry proteomics data have been deposited to the ProteomeXchange Consortium (http://proteomecentral.proteomexchange.org) via the PRIDE partner repository (77) with the dataset identifier PXD062352. Reviewers can access this repository via the username ‘’ and password ‘tOUIuuHykhtc’.

## Supplemental data

This article contains supplemental data:

**Table S1**. Set collision energy scheme for timsTOF SCP acquisition of MHC-I and MHC-II peptides.

**Table S2**. Overview of plate- and microfluidics-based immunopeptidomics approaches using immunopurification.

**Figure S1**. TimsTOF SCP polygons used for MHC-I and MHC-II peptide acquisition.

**Figure S2**. Effect of lysate volume and concentration on IP efficiency.

**Figure S3**. Optimization of MHC-I and MHC-II IP methods.

**Figure S4**. Optimization of the IP elution conditions.

**Figure S5**. Optimization of the peptide purification method.

**Figure S6**. Identified peptide length histograms of MHC-II pulldowns testing different antibodies and without prior MHC-I pulldown.

**Figure S7**. JY MHC class I and II predicted binding strengths.

**Figure S8**. Immunopeptidomics on HeLa and U937 cell dilutions.

**Figure S9**. Peptide length and Gibbs clusters of ultrasensitive JY cell dilutions.

**Figure S10**. Ultrasensitive immunopeptidomics in presence of n-dodecyl-β-D-maltoside.

**Figure S11**. Predicted binding strength for identified immunopeptides in the ultrasensitive JY dilution series.

**Figure S12**. Predicted non-binders and contaminant proteins increase with decreasing cell amount inputs.

**Figure S13**. Immunopeptidomics quality control of U937 cell cultures infected by *Listeria monocytogenes* and *Mycobacterium bovis* BCG.

**Figure S14**. Quantitative reproducibility and variation between BCG-infected and uninfected samples.

**Data S1**. Identified MHC-I immunopeptide sequences in JY serial cell dilutions [XLSX].

**Data S2**. Identified MHC-II immunopeptide sequences in JY serial cell dilutions [XLSX].

**Data S3**. Identified MHC-I immunopeptide sequences in HeLa serial cell dilutions [XLSX].

**Data S4**. Identified MHC-I immunopeptide sequences in U937 serial cell dilutions [XLSX].

**Data S5**. Identified MHC-I immunopeptide sequences in JY low input serial cell dilutions without the nonionic surfactant n-dodecyl-β-D-maltoside (DDM) [XLSX].

**Data S6**. Identified MHC-I immunopeptide sequences in JY low input serial cell dilutions with the nonionic surfactant n-dodecyl-β-D-maltoside (DDM) [XLSX].

**Data S7**. FragPipe HLA peptidome quantitative analysis of the JY low input serial dilutions without the nonionic surfactant n-dodecyl-β-D-maltoside (DDM) [XLSX].

**Data S8**. Identified human self-peptides and *Listeria* peptides during infection in U937 cells [XLSX].

**Data S9**. Identified human self-peptides and BCG peptides during infection in U937 cells [XLSX].

**Data S10**. Differential peptide abundance analysis during BCG infection [XLSX].

## Acknowledgements

We thank the VIB Proteomics Core for LC-MS/MS analysis. F.I., C.M., B.V. and I.L. acknowledge support from the Horizon Europe Project BAXERNA 2.0 [101080544]. F.I., I.L. and B.V. acknowledge support from Ghent University Concerted Research Action grant BOF21/GOA/033 and F.I. from Starting Grant BOF/STA/202209/011, as well as from the European Research Council (ERC Consolidator Grant #101089193). A.G. is supported by a PhD fellowship from the Higher Education Commission (HEC) Pakistan. This work was supported by the Research Foundation–Flanders (FWO) through a Junior Postdoctoral fellowship to F.T. (12AN524N) and a PhD fellowship for Strategic Basic Research to I.A. (1S40923N).

## Author contributions

**Investigation**: AG, LVM, FT, AS, KCF, LPC, KB, IA; **Formal analysis and data curation**: AG, LVM, FT, AS, PW; **Visualization**: AG, LVM, PW; **Resources**: BV, CD, FI; **Writing – Original Draft**: AG, FT, LVM, PW, FI; **Writing – Review & Editing**: AG, LVM, FT, PW, AS, BV, IL, IA, CD, SD, FI; **Conceptualization**: FI; **Supervision**: FT, FI. All authors read and approved the final manuscript.

## References

1. Pishesha, N., Harmand, T. J., and Ploegh, H. L. (2022) A guide to antigen processing and presentation. Nat Rev Immunol 22, 751–764

2. Thibault, P., and Perreault, C. (2022) Immunopeptidomics: Reading the Immune Signal That Defines Self From Nonself. Mol Cell Proteomics 21, 100234

3. Bassani-Sternberg, M., Pletscher-Frankild, S., Jensen, L. J., and Mann, M. (2015) Mass spectrometry of human leukocyte antigen class I peptidomes reveals strong effects of protein abundance and turnover on antigen presentation. Mol Cell Proteomics 14, 658–673

4. Caron, E., Vincent, K., Fortier, M. H., Laverdure, J. P., Bramoulle, A., Hardy, M. P., Voisin, G., Roux, P. P., Lemieux, S., Thibault, P., and Perreault, C. (2011) The MHC I immunopeptidome conveys to the cell surface an integrative view of cellular regulation. Mol Syst Biol 7, 533

5. Shapiro, I. E., and Bassani-Sternberg, M. (2023) The impact of immunopeptidomics: From basic research to clinical implementation. Semin Immunol 66, 101727

6. Gfeller, D., and Bassani-Sternberg, M. (2018) Predicting Antigen Presentation-What Could We Learn From a Million Peptides? Front Immunol 9, 1716

7. Flender, D., Vilenne, F., Adams, C., Boonen, K., Valkenborg, D., and Baggerman, G. (2025) Exploring the dynamic landscape of immunopeptidomics: Unravelling posttranslational modifications and navigating bioinformatics terrain. Mass Spectrom Rev 44, 599–629

8. Mayer, R. L., Verbeke, R., Asselman, C., Aernout, I., Gul, A., Eggermont, D., Boucher, K., Thery, F., Maia, T. M., Demol, H., Gabriels, R., Martens, L., Becavin, C., De Smedt, S. C., Vandekerckhove, B., Lentacker, I., and Impens, F. (2022) Immunopeptidomics-based design of mRNA vaccine formulations against Listeria monocytogenes. Nat Commun 13, 6075

9. Bettencourt, P., Muller, J., Nicastri, A., Cantillon, D., Madhavan, M., Charles, P. D., Fotso, C. B., Wittenberg, R., Bull, N., Pinpathomrat, N., Waddell, S. J., Stylianou, E., Hill, A. V. S., Ternette, N., and McShane, H. (2020) Identification of antigens presented by MHC for vaccines against tuberculosis. NPJ Vaccines 5, 2

10. Almujri, S. S., Stylianou, E., Nicastri, A., Satti, I., Korompis, M., Li, S., De Voss, C. J., Polo Peralta Alvarez, M., Tanner, R., Bettencourt, P. J. G., Ternette, N., and McShane, H. (2025) MetE: a promising protective antigen for tuberculosis vaccine development. Front Immunol 16, 1593263

11. Leddy, O., Ogongo, P., Huffaker, J., Gan, M., Milligan, R., Mahmud, S., Ni, H. M., Yuki, Y., Bobosha, K., Wassie, L., Carrington, M., Liu, Q., Ernst, J. D., White, F. M., and Bryson, B. D. (2025) Immunopeptidomics can inform the design of mRNA vaccines for the delivery of Mycobacterium tuberculosis MHC class II antigens. Sci Transl Med 17, eadw9184

12. Leddy, O., White, F. M., and Bryson, B. D. (2023) Immunopeptidomics reveals determinants of Mycobacterium tuberculosis antigen presentation on MHC class I. Elife 12

13. Leddy, O., Yuki, Y., Carrington, M., Bryson, B. D., and White, F. M. (2025) Targeting infection-specific peptides in immunopeptidomics studies for vaccine target discovery. J Exp Med 222

14. Cormican, J. A., Medfai, L., Wawrzyniuk, M., Pasen, M., Afrache, H., Fourny, C., Khan, S., Gneisse, P., Soh, W. T., Timelli, A., Nolfi, E., Pannekoek, Y., Cope, A., Urlaub, H., Sijts, A., Mishto, M., and Liepe, J. (2025) PEPSeek-Mediated Identification of Novel Epitopes From Viral and Bacterial Pathogens and the Impact on Host Cell Immunopeptidomes. Mol Cell Proteomics 24, 100937

15. Gul, A., Pewe, L. L., Willems, P., Mayer, R., Thery, F., Asselman, C., Aernout, I., Verbeke, R., Eggermont, D., Van Moortel, L., Upton, E., Zhang, Y., Boucher, K., Miret-Casals, L., Demol, H., De Smedt, S. C., Lentacker, I., Radoshevich, L., Harty, J. T., and Impens, F. (2024) Immunopeptidomics Mapping of Listeria monocytogenes T Cell Epitopes in Mice. Mol Cell Proteomics 23, 100829

16. Karunakaran, K. P., Yu, H., Jiang, X., Chan, Q., Goldberg, M. F., Jenkins, M. K., Foster, L. J., and Brunham, R. C. (2017) Identification of MHC-Bound Peptides from Dendritic Cells Infected with Salmonella enterica Strain SL1344: Implications for a Nontyphoidal Salmonella Vaccine. J Proteome Res 16, 298–306

17. Zhang, B., and Bassani-Sternberg, M. (2023) Current perspectives on mass spectrometry-based immunopeptidomics: the computational angle to tumor antigen discovery. J Immunother Cancer 11

18. Bassani-Sternberg, M., Braunlein, E., Klar, R., Engleitner, T., Sinitcyn, P., Audehm, S., Straub, M., Weber, J., Slotta-Huspenina, J., Specht, K., Martignoni, M. E., Werner, A., Hein, R., D, H. B., Peschel, C., Rad, R., Cox, J., Mann, M., and Krackhardt, A. M. (2016) Direct identification of clinically relevant neoepitopes presented on native human melanoma tissue by mass spectrometry. Nat Commun 7, 13404

19. Shapiro, I. E., Huber, F., Michaux, J., and Bassani-Sternberg, M. (2025) Sensitive neoantigen discovery by real-time mutanome-guided immunopeptidomics. Nat Commun 16, 7269

20. Apavaloaei, A., Zhao, Q., Hesnard, L., Cahuzac, M., Durette, C., Larouche, J. D., Hardy, M. P., Vincent, K., Brochu, S., Laverdure, J. P., Lanoix, J., Courcelles, M., Gendron, P., Lajoie, M., Ruiz Cuevas, M. V., Kina, E., Perrault, J., Humeau, J., Ehx, G., Lemieux, S., Watson, I. R., Speiser, D. E., Bassani-Sternberg, M., Thibault, P., and Perreault, C. (2025) Tumor antigens preferentially derive from unmutated genomic sequences in melanoma and non-small cell lung cancer. Nat Cancer 6, 1419–1437

21. Ely, Z. A., Kulstad, Z. J., Gunaydin, G., Addepalli, S., Verzani, E. K., Casarrubios, M., Clauser, K. R., Wang, X., Lippincott, I. E., Louvet, C., Schmitt, T., Kapner, K. S., Agus, M. P., Hennessey, C. J., Cleary, J. M., Hadrup, S. R., Klaeger, S., Su, J., Jaeger, M., Wolpin, B. M., Raghavan, S., Smith, E. L., Greenberg, P. D., Aguirre, A. J., Abelin, J. G., Carr, S. A., Jacks, T., and Freed-Pastor, W. A. (2025) Pancreatic cancer-restricted cryptic antigens are targets for T cell recognition. Science 388, eadk3487

22. Purcell, A. W., Ramarathinam, S. H., and Ternette, N. (2019) Mass spectrometry-based identification of MHC-bound peptides for immunopeptidomics. Nat Protoc 14, 1687–1707

23. Marino, F., Chong, C., Michaux, J., and Bassani-Sternberg, M. (2019) High-Throughput, Fast, and Sensitive Immunopeptidomics Sample Processing for Mass Spectrometry. Methods Mol Biol 1913, 67–79

24. Mohan, S. V., Datta, K. K., Ziegman, R., Smith, C., and Gowda, H. (2021) Protocol for purification and identification of MHC class I immunopeptidome from cancer cell lines. STAR Protoc 2, 100385

25. Kowalewski, D. J., and Stevanovic, S. (2013) Biochemical large-scale identification of MHC class I ligands. Methods Mol Biol 960, 145–157

26. Chong, C., Marino, F., Pak, H., Racle, J., Daniel, R. T., Muller, M., Gfeller, D., Coukos, G., and Bassani-Sternberg, M. (2018) High-throughput and Sensitive Immunopeptidomics Platform Reveals Profound Interferongamma-Mediated Remodeling of the Human Leukocyte Antigen (HLA) Ligandome. Mol Cell Proteomics 17, 533–548

27. Wacker, M., Bauer, J., Wessling, L., Dubbelaar, M., Nelde, A., Rammensee, H. G., and Walz, J. S. (2023) Immunoprecipitation methods impact the peptide repertoire in immunopeptidomics. Front Immunol 14, 1219720

28. Phulphagar, K. M., Ctortecka, C., Jacome, A. S. V., Klaeger, S., Verzani, E. K., Hernandez, G. M., Udeshi, N. D., Clauser, K. R., Abelin, J. G., and Carr, S. A. (2023) Sensitive, High-Throughput HLA-I and HLA-II Immunopeptidomics Using Parallel Accumulation-Serial Fragmentation Mass Spectrometry. Mol Cell Proteomics 22, 100563

29. Beyrle, J., Distler, U., Gomez-Zepeda, D., and Tenzer, S. (2024) MAETi: Mild acid elution in a tip enables immunopeptidome profiling from 25,000 cells. bioRxiv, 2024.2012.2020.628848

30. Sturm, T., Sautter, B., Worner, T. P., Stevanovic, S., Rammensee, H. G., Planz, O., Heck, A. J. R., and Aebersold, R. (2021) Mild Acid Elution and MHC Immunoaffinity Chromatography Reveal Similar Albeit Not Identical Profiles of the HLA Class I Immunopeptidome. J Proteome Res 20, 289–304

31. Bathini, M., Bocaniciu, D., Johnson, F. D., de Jong, R. C. P., Yu, F., Aloi, V. D., Kuiken, M. C., Mors, J. R., Giebel, L., Champagne, J., Bleijerveld, O., Agami, R., Dijkstra, K. K., Thommen, D. S., Nesvizhskii, A. I., and Lindeboom, R. G. H. (2025) MHC1-TIP enables single-tube multimodal immunopeptidome profiling and uncovers intratumoral heterogeneity in antigen presentation. bioRxiv, 2025.2007.2017.664894

32. Gomez-Zepeda, D., Arnold-Schild, D., Beyrle, J., Declercq, A., Gabriels, R., Kumm, E., Preikschat, A., Lacki, M. K., Hirschler, A., Rijal, J. B., Carapito, C., Martens, L., Distler, U., Schild, H., and Tenzer, S. (2024) Thunder-DDA-PASEF enables high-coverage immunopeptidomics and is boosted by MS(2)Rescore with MS(2)PIP timsTOF fragmentation prediction model. Nat Commun 15, 2288

33. Willems, P., Thery, F., Van Moortel, L., De Meyer, M., Staes, A., Gul, A., Kovalchuke, L., Declercq, A., Devreese, R., Bouwmeester, R., Gabriels, R., Martens, L., and Impens, F. (2025) Maximizing Immunopeptidomics-Based Bacterial Epitope Discovery by Multiple Search Engines and Rescoring. J Proteome Res 24, 2141–2151

34. Willems, P., Staes, A., Miret-Casals, L., Demichev, V., Devos, S., and Impens, F. (2025) Data-Independent Immunopeptidomics Discovery of Low-Abundant Bacterial Epitopes. J Proteome Res

35. Pak, H., Michaux, J., Huber, F., Chong, C., Stevenson, B. J., Muller, M., Coukos, G., and Bassani-Sternberg, M. (2021) Sensitive Immunopeptidomics by Leveraging Available Large-Scale Multi-HLA Spectral Libraries, Data-Independent Acquisition, and MS/MS Prediction. Mol Cell Proteomics 20, 100080

36. Wahle, M., Thielert, M., Zwiebel, M., Skowronek, P., Zeng, W. F., and Mann, M. (2024) IMBAS-MS Discovers Organ-Specific HLA Peptide Patterns in Plasma. Mol Cell Proteomics 23, 100689

37. Declercq, A., Bouwmeester, R., Hirschler, A., Carapito, C., Degroeve, S., Martens, L., and Gabriels, R. (2022) MS(2)Rescore: Data-Driven Rescoring Dramatically Boosts Immunopeptide Identification Rates. Mol Cell Proteomics 21, 100266

38. Yang, K. L., Yu, F., Teo, G. C., Li, K., Demichev, V., Ralser, M., and Nesvizhskii, A. I. (2023) MSBooster: improving peptide identification rates using deep learning-based features. Nat Commun 14, 4539

39. Hunt, D. F., Henderson, R. A., Shabanowitz, J., Sakaguchi, K., Michel, H., Sevilir, N., Cox, A. L., Appella, E., and Engelhard, V. H. (1992) Characterization of peptides bound to the class I MHC molecule HLA-A2.1 by mass spectrometry. Science 255, 1261–1263

40. Zhang, L., McAlpine, P. L., Heberling, M. L., and Elias, J. E. (2021) Automated Ligand Purification Platform Accelerates Immunopeptidome Analysis by Mass Spectrometry. J Proteome Res 20, 393–408

41. Pollock, S. B., Rose, C. M., Darwish, M., Bouziat, R., Delamarre, L., Blanchette, C., and Lill, J. R. (2021) Sensitive and Quantitative Detection of MHC-I Displayed Neoepitopes Using a Semiautomated Workflow and TOMAHAQ Mass Spectrometry. Mol Cell Proteomics 20, 100108

42. Lim Kam Sian, T. C. C., Goncalves, G., Steele, J. R., Shamekhi, T., Bramberger, L., Jin, D., Shahbazy, M., Purcell, A. W., Ramarathinam, S., Stoychev, S., and Faridi, P. (2023) SAPrIm, a semi-automated protocol for mid-throughput immunopeptidomics. Front Immunol 14, 1107576

43. Li, X., Pak, H. S., Huber, F., Michaux, J., Taillandier-Coindard, M., Altimiras, E. R., and Bassani-Sternberg, M. (2023) A microfluidics-enabled automated workflow of sample preparation for MS-based immunopeptidomics. Cell Rep Methods 3, 100479

44. Abelin, J. G., Bergstrom, E. J., Rivera, K. D., Taylor, H. B., Klaeger, S., Xu, C., Verzani, E. K., Jackson White, C., Woldemichael, H. B., Virshup, M., Olive, M. E., Maynard, M., Vartany, S. A., Allen, J. D., Phulphagar, K., Harry Kane, M., Rachimi, S., Mani, D. R., Gillette, M. A., Satpathy, S., Clauser, K. R., Udeshi, N. D., and Carr, S. A. (2023) Workflow enabling deepscale immunopeptidome, proteome, ubiquitylome, phosphoproteome, and acetylome analyses of sample-limited tissues. Nat Commun 14, 1851

45. Feola, S., Haapala, M., Peltonen, K., Capasso, C., Martins, B., Antignani, G., Federico, A., Pietiainen, V., Chiaro, J., Feodoroff, M., Russo, S., Rannikko, A., Fusciello, M., Koskela, S., Partanen, J., Hamdan, F., Tahka, S. M., Ylosmaki, E., Greco, D., Gronholm, M., Kekarainen, T., Eshaghi, M., Gurvich, O. L., Yla-Herttuala, S., RM, M. B., Lehtio, J., Sikanen, T. M., and Cerullo, V. (2021) PeptiCHIP: A Microfluidic Platform for Tumor Antigen Landscape Identification. ACS Nano 15, 15992–16010

46. Tanuwidjaya, E., Lim Kam Sian, T. C. C., Steele, J. R., Goncalves, G., Woodhouse, I. B., Chang, J., Ooi, J. D., Schittenhelm, R. B., and Faridi, P. (2025) SAPrIm 2.0: a semi-automated protocol for mid-throughput soluble HLA immunopeptidomics. Front Immunol 16, 1546629

47. Rijkers, G. T., Roord, J. J., Koning, F., Kuis, W., and Zegers, B. J. (1987) Phenotypical and functional analysis of B lymphocytes of two siblings with combined immunodeficiency and defective expression of major histocompatibility complex (MHC) class II antigens on mononuclear cells. J Clin Immunol 7, 98–106

48. Kong, A. T., Leprevost, F. V., Avtonomov, D. M., Mellacheruvu, D., and Nesvizhskii, A. I. (2017) MSFragger: ultrafast and comprehensive peptide identification in mass spectrometry-based proteomics. Nat Methods 14, 513–520

49. Eng, J. K., Jahan, T. A., and Hoopmann, M. R. (2013) Comet: an open-source MS/MS sequence database search tool. Proteomics 13, 22–24

50. Lazear, M. R. (2023) Sage: An Open-Source Tool for Fast Proteomics Searching and Quantification at Scale. J Proteome Res 22, 3652–3659

51. Zhang, J., Xin, L., Shan, B., Chen, W., Xie, M., Yuen, D., Zhang, W., Zhang, Z., Lajoie, G. A., and Ma, B. (2012) PEAKS DB: de novo sequencing assisted database search for sensitive and accurate peptide identification. Mol Cell Proteomics 11, M111 010587

52. Declercq, A., Devreese, R., Scheid, J., Jachmann, C., Van Den Bossche, T., Preikschat, A., Gomez-Zepeda, D., Rijal, J. B., Hirschler, A., Krieger, J. R., Srikumar, T., Rosenberger, G., Martelli, C., Trede, D., Carapito, C., Tenzer, S., Walz, J. S., Degroeve, S., Bouwmeester, R., Martens, L., and Gabriels, R. (2025) TIMS(2)Rescore: A Data Dependent Acquisition-Parallel Accumulation and Serial Fragmentation-Optimized Data-Driven Rescoring Pipeline Based on MS(2)Rescore. J Proteome Res 24, 1067–1076

53. Reynisson, B., Alvarez, B., Paul, S., Peters, B., and Nielsen, M. (2020) NetMHCpan-4.1 and NetMHCIIpan-4.0: improved predictions of MHC antigen presentation by concurrent motif deconvolution and integration of MS MHC eluted ligand data. Nucleic Acids Res 48, W449–W454

54. Nilsson, J. B., Kaabinejadian, S., Yari, H., Kester, M. G. D., van Balen, P., Hildebrand, W. H., and Nielsen, M. (2023) Accurate prediction of HLA class II antigen presentation across all loci using tailored data acquisition and refined machine learning. Sci Adv 9, eadj6367

55. Andreatta, M., Alvarez, B., and Nielsen, M. (2017) GibbsCluster: unsupervised clustering and alignment of peptide sequences. Nucleic Acids Res 45, W458–W463

56. Yu, F., Haynes, S. E., and Nesvizhskii, A. I. (2021) IonQuant Enables Accurate and Sensitive Label-Free Quantification With FDR-Controlled Match-Between-Runs. Mol Cell Proteomics 20, 100077

57. Smyth, G. K. (2004) Linear models and empirical bayes methods for assessing differential expression in microarray experiments. Stat Appl Genet Mol Biol 3, Article3

58. van Ooijen, M. P., Jong, V. L., Eijkemans, M. J. C., Heck, A. J. R., Andeweg, A. C., Binai, N. A., and van den Ham, H. J. (2018) Identification of differentially expressed peptides in high-throughput proteomics data. Brief Bioinform 19, 971–981

59. Raudvere, U., Kolberg, L., Kuzmin, I., Arak, T., Adler, P., Peterson, H., and Vilo, J. (2019) g:Profiler: a web server for functional enrichment analysis and conversions of gene lists (2019 update). Nucleic Acids Res 47, W191–W198

60. Hunter, J. D. (2007) Matplotlib: A 2D Graphics Environment. Computing in Science & Engineering 9, 90–95

61. Waskom, M. L. (2021) seaborn: statistical data visualization. Journal of Open Source Software 6, 3021

62. Tareen, A., and Kinney, J. B. (2020) Logomaker: beautiful sequence logos in Python. Bioinformatics 36, 2272–2274

63. Lex, A., Gehlenborg, N., Strobelt, H., Vuillemot, R., and Pfister, H. (2014) UpSet: Visualization of Intersecting Sets. IEEE Trans Vis Comput Graph 20, 1983–1992

64. Guasp, P., Alvarez-Navarro, C., Gomez-Molina, P., Martin-Esteban, A., Marcilla, M., Barnea, E., Admon, A., and Lopez de Castro, J. A. (2016) The Peptidome of Behcet’s Disease-Associated HLA-B*51:01 Includes Two Subpeptidomes Differentially Shaped by Endoplasmic Reticulum Aminopeptidase 1. Arthritis Rheumatol 68, 505–515

65. Tsai, C. F., Zhang, P., Scholten, D., Martin, K., Wang, Y. T., Zhao, R., Chrisler, W., Patel, D. B., Dou, M., Jia, Y., Reduzzi, C., Liu, X., Moore, R. J., Burnum-Johnson, K. E., Lin, M. H., Hsu, C. C., Jacobs, J. M., Kagan, J., Srivastava, S., Rodland, K. D., Steven Wiley, H., Qian, W. J., Smith, R. D., Zhu, Y., Cristofanilli, M., Liu, T., Liu, H., and Shi, T. (2021) Surfactant-assisted one-pot sample preparation for label-free single-cell proteomics. Commun Biol 4, 265

66. Aernout, I., Verbeke, R., Thery, F., Willems, P., Elia, U., De Smedt, S. C., Rappuoli, R., Peer, D., Impens, F., and Lentacker, I. (2025) Challenges and opportunities in mRNA vaccine development against bacteria. Nat Microbiol 10, 1816–1828

67. Karunakaran, K. P., Yu, H., Jiang, X., Chan, Q. W. T., Sigola, L., Millis, L. A., Chen, J., Tang, P., Foster, L. J., and Brunham, R. C. (2024) Immunoproteomic discovery of Mycobacterium bovis antigens, including the surface lipoprotein Mpt83 as a T cell antigen useful for vaccine development. Vaccine 42, 126266

68. Flyer, D. C., Ramakrishna, V., Miller, C., Myers, H., McDaniel, M., Root, K., Flournoy, C., Engelhard, V. H., Canaday, D. H., Marto, J. A., Ross, M. M., Hunt, D. F., Shabanowitz, J., and White, F. M. (2002) Identification by mass spectrometry of CD8(+)-T-cell Mycobacterium tuberculosis epitopes within the Rv0341 gene product. Infect Immun 70, 2926–2932

69. Cooper, A. M., Mayer-Barber, K. D., and Sher, A. (2011) Role of innate cytokines in mycobacterial infection. Mucosal Immunol 4, 252–260

70. Ford, S. G., Caswell, P., Brough, D., and Seoane, P. I. (2025) The secretion of interleukin-1beta. Cytokine Growth Factor Rev 84, 101–113

71. Monteleone, M., Stow, J. L., and Schroder, K. (2015) Mechanisms of unconventional secretion of IL-1 family cytokines. Cytokine 74, 213–218

72. Oliinyk, D., Gurung, H. R., Zhou, Z., Leskoske, K., Rose, C. M., and Klaeger, S. (2025) diaPASEF Analysis for HLA-I Peptides Enables Quantification of Common Cancer Neoantigens. Mol Cell Proteomics 24, 100938

73. Sinn, L. R., Szyrwiel, L., Grossmann, J., Lau, K., Faisst, K., Qin, D., Mutschler, F., Khoury, L., Leduc, A., Ralser, M., Coscia, F., Selbach, M., Slavov, N., Nagaraj, N., Steger, M., and Demichev, V. (2025) Slice-PASEF: Maximising Ion Utilisation in LC-MS Proteomics. bioRxiv, 2022.2010.2031.514544

74. Skowronek, P., Krohs, F., Lubeck, M., Wallmann, G., Itang, E. C. M., Koval, P., Wahle, M., Thielert, M., Meier, F., Willems, S., Raether, O., and Mann, M. (2023) Synchro-PASEF Allows Precursor-Specific Fragment Ion Extraction and Interference Removal in Data-Independent Acquisition. Mol Cell Proteomics 22, 100489

75. Lanoix, J., Durette, C., Courcelles, M., Cossette, E., Comtois-Marotte, S., Hardy, M. P., Cote, C., Perreault, C., and Thibault, P. (2018) Comparison of the MHC I Immunopeptidome Repertoire of B-Cell Lymphoblasts Using Two Isolation Methods. Proteomics 18, e1700251

76. Schaefer, Z., Iradukunda, J., Lumngwena, E. N., Basso, K. B., Blackburn, J. M., and Parker, I. K. (2024) Multilevel Proteomics Reveals Epigenetic Signatures in BCG-Mediated Macrophage Activation. Mol Cell Proteomics 23, 100851

77 Perez-Riverol, Y., Bandla, C., Kundu, D. J., Kamatchinathan, S., Bai, J., Hewapathirana, S., John, N. S., Prakash, A., Walzer, M., Wang, S., and Vizcaino, J. (2025) The PRIDE database at 20 years: 2025 update. Nucleic Acids Res 53, D543–D553

