## Supplemental Data for "A platform for high-throughput and ultrasensitive immunopeptidomics"

**Author affiliations:**

**TABLE OF CONTENTS**


| | |
|---|---|
| Supplemental Table S1 | Set collision energy scheme for timsTOF SCP acquisition of MHC-I and MHC-II peptides. |
| Supplemental Table S2 | Overview of plate- and microfluidics-based immunopeptidomics approaches using immunopurification. |
| Supplemental Figure S1 | TimsTOF SCP polygons used for MHC-I and MHC-II peptide acquisition. |
| Supplemental Figure S2 | Effect of lysate volume and concentration on IP efficiency. |
| Supplemental Figure S3 | Optimization of MHC-I and MHC-II IP methods. |
| Supplemental Figure S4 | Optimization of the IP elution conditions. |
| Supplemental Figure S5 | Optimization of the peptide purification method. |
| Supplemental Figure S6 | Identified peptide length histograms of MHC-II pulldowns testing different antibodies and without prior MHC-I pulldown. |
| Supplemental Figure S7 | JY MHC class I and II predicted binding strengths. |
| Supplemental Figure S8 | Immunopeptidomics on HeLa and U937 cell dilutions. |
| Supplemental Figure S9 | Peptide length and Gibbs clusters of ultrasensitive JY cell dilutions. |
| Supplemental Figure S10 | Ultrasensitive immunopeptidomics in presence of n-dodecyl-β-D-maltoside. |
| Supplemental Figure S11 | Predicted binding strength for identified immunopeptides in the ultrasensitive JY dilution series. |
| Supplemental Figure S12 | Predicted non-binders and contaminant proteins increase with decreasing cell amount inputs. |
| Supplemental Figure S13 | Immunopeptidomics quality control of U937 cell cultures infected by *Listeria monocytogenes* and *Mycobacterium bovis* BCG. |
| Supplemental Figure S14 | Quantitative reproducibility and variation between BCG-infected and uninfected samples. |

| | |
|---|---|
| Supplemental Data S1 | Identified MHC-I immunopeptide sequences in JY serial cell dilutions [XLSX]. |
| Supplemental Data S2 | Identified MHC-II immunopeptide sequences in JY serial cell dilutions [XLSX]. |
| Supplemental Data S3 | Identified MHC-I immunopeptide sequences in HeLa serial cell dilutions [XLSX]. |
| Supplemental Data S4 | Identified MHC-I immunopeptide sequences in U937 serial cell dilutions [XLSX]. |
| Supplemental Data S5 | Identified MHC-I immunopeptide sequences in JY low input serial cell dilutions without the nonionic surfactant n-dodecyl-β-D-maltoside (DDM) [XLSX]. |
| Supplemental Data S6 | Identified MHC-I immunopeptide sequences in JY low input serial cell dilutions with the nonionic surfactant n-dodecyl-β-D-maltoside (DDM) [XLSX]. |
| Supplemental Data S7 | FragPipe HLA peptidome quantitative analysis of the JY low input serial dilutions without the nonionic surfactant n-dodecyl-β-D-maltoside (DDM) [XLSX]. |
| Supplemental Data S8 | Identified human self-peptides and *Listeria* peptides during infection in U937 cells [XLSX]. |
| Supplemental Data S9 | Identified human self-peptides and BCG peptides during infection in U937 cells [XLSX]. |
| Supplemental Data S10 | Differential peptide abundance analysis during BCG infection [XLSX]. |

#### Supplementary Tables

**Supplementary Table S1.** Set collision energy scheme for timsTOF SCP acquisition of MHC-I and MHC-II peptides.

| $1/K_0$ [Vs cm$^{-2}$] | Collision energy [eV] |
|---|---|
| 0.7 | 20 |
| 1.06 | 30 |
| 1.16 | 40 |
| 1.34 | 40 |
| 1.68 | 70 |

**Supplementary Table S2.** Overview of plate- and microfluidics-based immunopeptidomics approaches using immunopurification. Details on sample preparation and data acquisition/analysis are provided. Peptides identified were reported at 1% FDR. Missing information is labeled as not specified (n.s.). Abbreviations: LAUD, lung adenocarcinoma; MHC-Ip, MHC class I peptides; MHC-IIp, MHC class II peptides; PDAC, pancreatic ductal adenocarcinoma; Prot-A, protein A.

| Study | Input tissue (g) or cells (number) | Sample preparation (lysis, IP, elution and purification) | | | | | | Data acquisition and analysis | |
|---|---|---|---|---|---|---|---|---|---|
| | | Antibody to bead ratio | Lysis volume | Automatization (# samples) | IP condition | MHCp elution | MHCp purification | MS instrument | Number of peptides identified (FDR 1%) |
| Chong et al. (26) | Meningioma tissue (1g) and 7 cell lines (100 MY) | 5 mg/mL Prot-A Sepharose | 10 mL (tissue), 1 mL (cells) | 96 well microplate 3 µm glass fiber and 10 µm polypropylene membranes (n=96) | Gravity flowthrough at 4°C | 1% TFA | Sep-Pak tC18 100 mg | QExactive HF | MHC-Ip: 3,293-13,696 MHC-IIp: 7,210-10,060 |
| Zhang et al. (40) | Raji cells (100 MY) | 250 µg/mL Prot-A cartridges (AssayMAP Bravo) | 0.8 mL | 96 well deep plate for lysate (n=96) | Flow rate 10 µL/min (80 min) | 10% acetic acid | 10 kDa spin column, AssayMAP's RP-S cartridge | LTQ Orbitrap Elite, Fusion Lumos | MHC-Ip: 5,578 MHC-IIp: 8,250 |
| Pollock et al. (41) | MC38 and GRANTA cells (250 MY) | 1mg/mL Prot-A cartridge | 5 mL | 96-well deep plate for lysate (n=96) | Flow rate 20 µL/min | 1% acetate | C18 Cartridges | Fusion Lumos Tribrid | MHC-Ip: > 4,000 |
| Abelin et al. (44) | LAUD tumor (50 mg) | 0.015 mg/37.5µL Gammabind Sepharose | 1.2 mL | IP in tubes and transferred to 10 µm PE fritted plate (n.s.) | Incubating tubes with beads and lysate on a rotor at 4 °C (3 h) | 10% acetic acid | Sep-Pak tC18 40 mg | Orbitrap Exploris 480 | MHC-Ip: 8,278-13,727 MHC-IIp: 1,123-9,726 |
| Lim Kam Sian et al. (42) | MDA-MB-231 cells (0.5 to 50 MY) | 0.5 mg/80 µL MagReSyn Prot-A | 300-600 µL | Kingfisher 96-well plate (n=12) | 1 h | 0.1% TFA | C-18 stage tips | Orbitrap Exploris 480 | MHC-Ip: ~ 400-3,000 |
| Phulphagar et al. (28) | Melanoma tumor (25 mg) and A375 and PDAC cell lines (1 to 40 MY) | 0.015 mg/37.5 µL Gammabind Sepharose | 1.2 mL | IP in 96-well deep plate and transferred to 10 µm PE fritted plate (n=96) | End-to-end rotation (3h) | 10% acetic acid | Sep-Pak tC18 40 mg | TimsTOF SCP | MHC-Ip: 7,000-15,000 MHC-IIp: up to 15,000 from 40e6 |
| Feola et al. (45) | JY cells (1 to 50 MY), ovarian tumor (10 to 60 mg), organoids | 45 µg biotinylated antibody | n.s. | IP on PeptiCHIP (n=1) | n.s. | 7% acetic acid in 50% meOH | SepPac-C18 cartridges | TimsTOF Pro | MHC-Ip: 8,00-5,590 from 1 to 50 MY JY cells |
| Li et al. (43) | Melanoma tumor (5 to 50 mg), RA957 cells (0.2 to 10 MY) | 3 mg/mL Prot-Sepharose | 100-200 µL | IP on Chip (n=12) | At lowest flow rate | 10% acetic acid | C-18 cartridges | Q Exactive HF-X | MHC-Ip: 4,000-15,000 |
| Tanuwidjaya et al. (46) | 500 µL plasma | 100 µg/100 µL MagReSyn Prot-A | n.s. | Kingfisher, 96-well plate (n=96) | 1 h | 10% acetic acid | SDB-XC stage tips | Orbitrap Exploris 480 | MHC-Ip: 1,257-4,226 from 100 µL to 1 mL plasma |

### Supplementary Figures

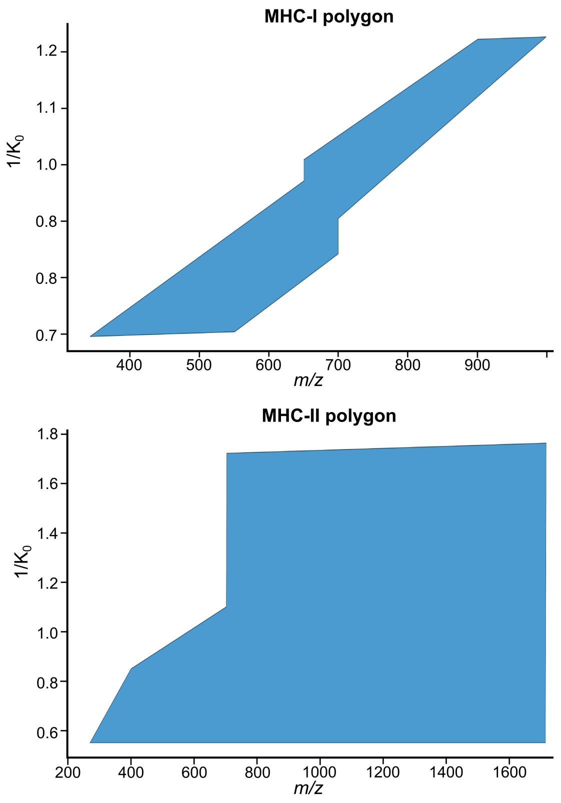


**Supplementary Figure S1**. **TimsTOF SCP polygons used for MHC-I and MHC-II peptide acquisition.** The blue area indicates the polygon across the inverse reduced ion mobility (1/$K_0$) vs *m/z* dimension. In case of MHC-I peptides (*Top*), a more restricted "Thunder" polygon (28, 32) is used, while a broad polygon for MHC-II peptides (*Bottom*). Polygon figures were extracted from method reports generated by the package timsCompare (v1.2) (https://github.com/kronigert/timsCompare).

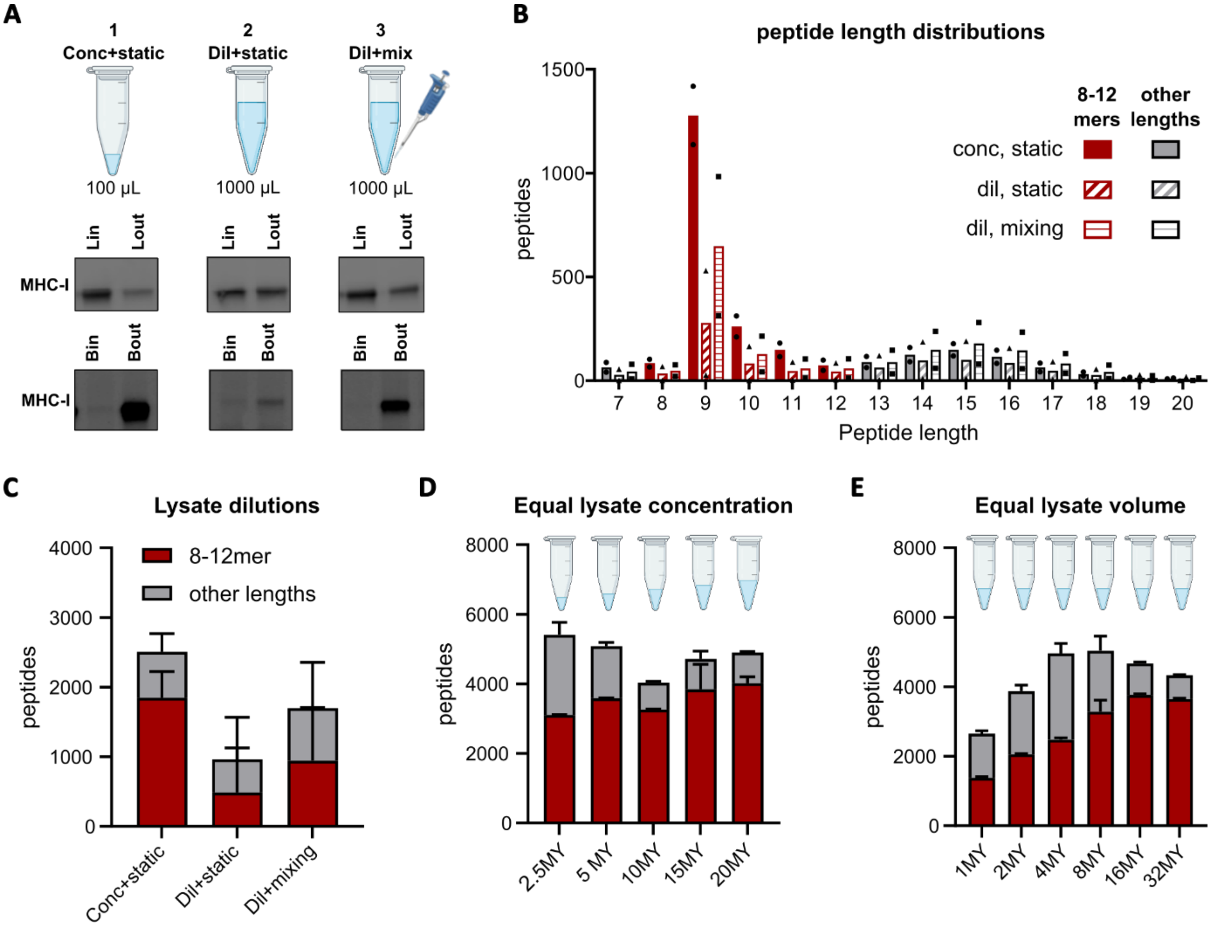


**Supplementary Figure S2**. **Effect of lysate volume and concentration on IP efficiency.** All data are representative of two technical replicates per condition. Washing and peptide clean-up steps were performed according to a standard immunopeptidomics workflow (8). Samples were analyzed on a timsTOF SCP instrument in-line coupled to an EvoSep One chromatography system (25% sample injection). Peptides were identified using PEAKS (v11.0) (51). **(A-C)** 10 million JY cells were lysed in 100 µL ("Conc") or 1000 µL ("Dil") lysis buffer and incubated for 1 h at 4°C with 300 µg W6/32 HLA class I antibody crosslinked to 100 µL packed protein A beads in 96-well plates, with or without mixing. **(A)** Control immunoblots incubated with antibody against HLA class I. Abbreviations: Lin, lysate before IP; Lout, lysate after IP; Bin, beads before IP; Bout, beads after IP. **(B)** Identified peptide length distribution for all three conditions. Data points indicate technical replicates. **(C)** Number of identified 8-12mer peptides and peptides of other lengths in all three conditions. **(D)** 2.5 to 20 million JY cells were lysed in 100 µL lysis buffer per 10 million cells and incubated for 1 hour at 4°C with 300 µg W6/32 antibody crosslinked to 100 µL packed protein A beads. The graph represents the number of identified 8-12mer peptides and peptides of other lengths identified per condition. **(E)** 1 to 32 million JY cells were lysed in 100 µL lysis buffer and incubated for 1 hour at 4°C with 300 µg W6/32 antibody crosslinked to 100 µL packed protein A beads. The graph represents the number of identified 8-12mer peptides and peptides of other lengths identified per condition. Figure created with BioRender.com.

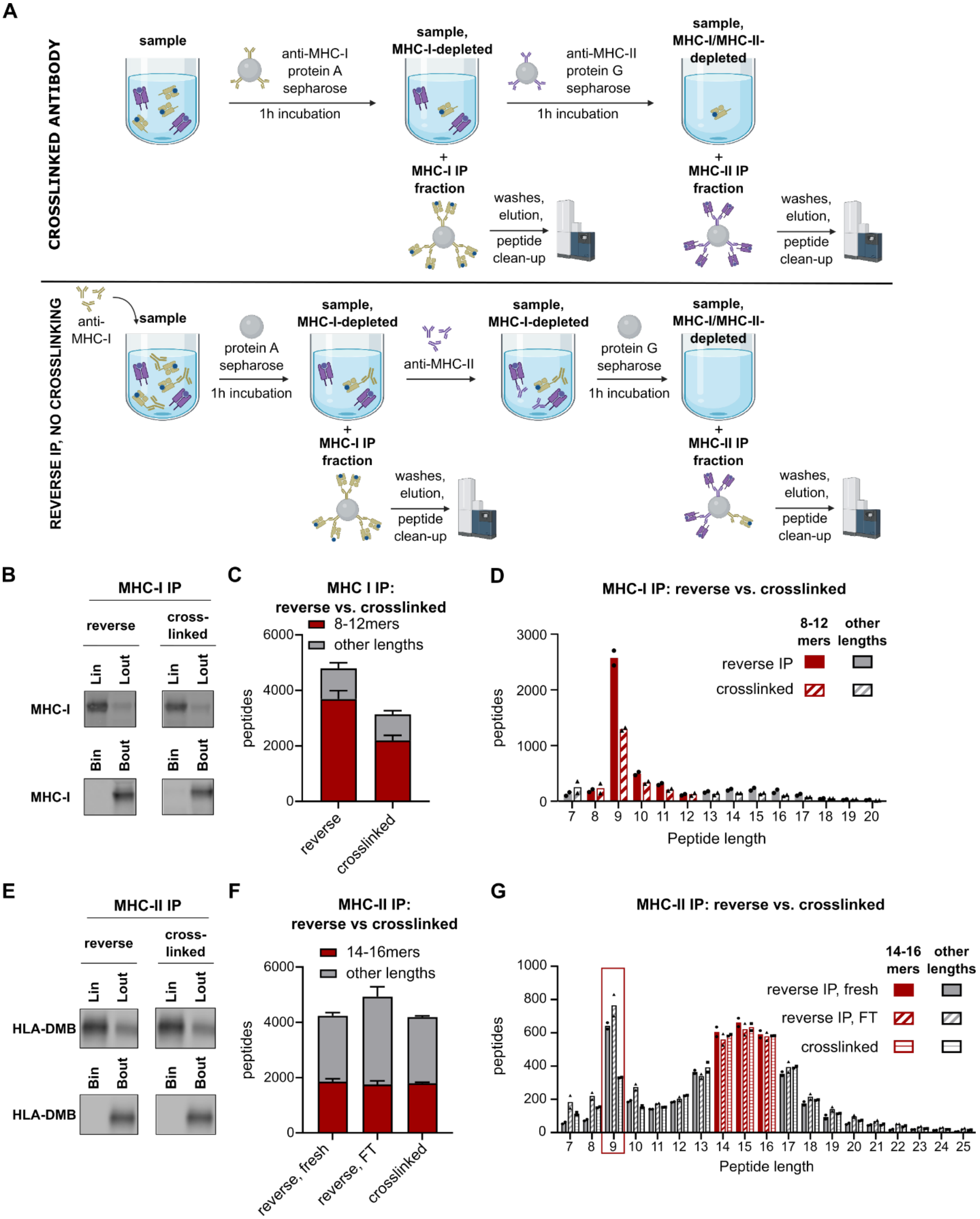


**Supplementary Figure S3. Optimization of MHC-I and MHC-II IP methods.** 10 million JY cells were lysed in 100 µL lysis buffer and subjected to various IP methods, followed by washing steps and peptide clean-up according to the standard immunopeptidomics workflow. Samples were run on a timsTOF SCP instrument in-coupled to an Evosep One chromatography system (25% sample injection). Data represent the average of two technical replicates. Peptides were identified using PEAKS (v11.0) (51). **(A)** Workflow overview for reverse IP and IP with crosslinked antibody. Figure created with Biorender. **(B-D)** Comparison of reverse IP and IP with crosslinked antibody for MHC class I. In the reverse IP, 100 µL lysate was mixed with 300 µg W6/32 antibody diluted in 100 µL 50 mM Tris pH7.5 150 mM NaCl before 1h incubation with 100 µL packed protein A beads (4°C). In the regular IP, 100 µL lysate was mixed with 100 µL 50 mM Tris pH7.5 150 mM NaCl and incubated with 300 µg W6/32 antibody crosslinked to 100 µL packed protein A beads for 1h at 4°C. **(B)** Control immunoblots, incubated with antibody against HLA class I. Lin, lysate before IP; Lout, lysate after IP; Bin, beads before IP; Bout, beads after IP. **(C)** Number of identified 8- to 12mer peptides (red) and peptides of other lengths (grey) in both conditions. **(D)** Peptide length distributions for both conditions. **(E-G)** Comparison of reverse IP and IP with crosslinked antibody for MHC class II. In the reverse IP, 175 µL fresh lysate ('reverse, fresh') or 175 µL lysate flow-through after MHC-I IP ('reverse, FT') was mixed with 150 µg PdV5.2 MHC class II antibody diluted in 125 µL 50 mM Tris pH7.5 150 mM NaCl before 1h incubation with 100 µL packed protein G beads (4°C). In the regular IP, 100 µL lysate was mixed with 100 µL 50 mM Tris pH7.5 150 mM NaCl and incubated with 150 µg PdV5.2 MHC class II antibody crosslinked to 100 µL packed protein G beads for 1h at 4°C. **(E)** Control immunoblots, incubated with antibody against HLA-DMB. Lin, lysate before IP; Lout, lysate after IP; Bin, beads before IP; Bout, beads after IP. **(F)** Number of identified 14- to 16-mer peptides (red) and peptides of other lengths (grey) identified in all conditions. **(G)** Peptide length distributions for all conditions.

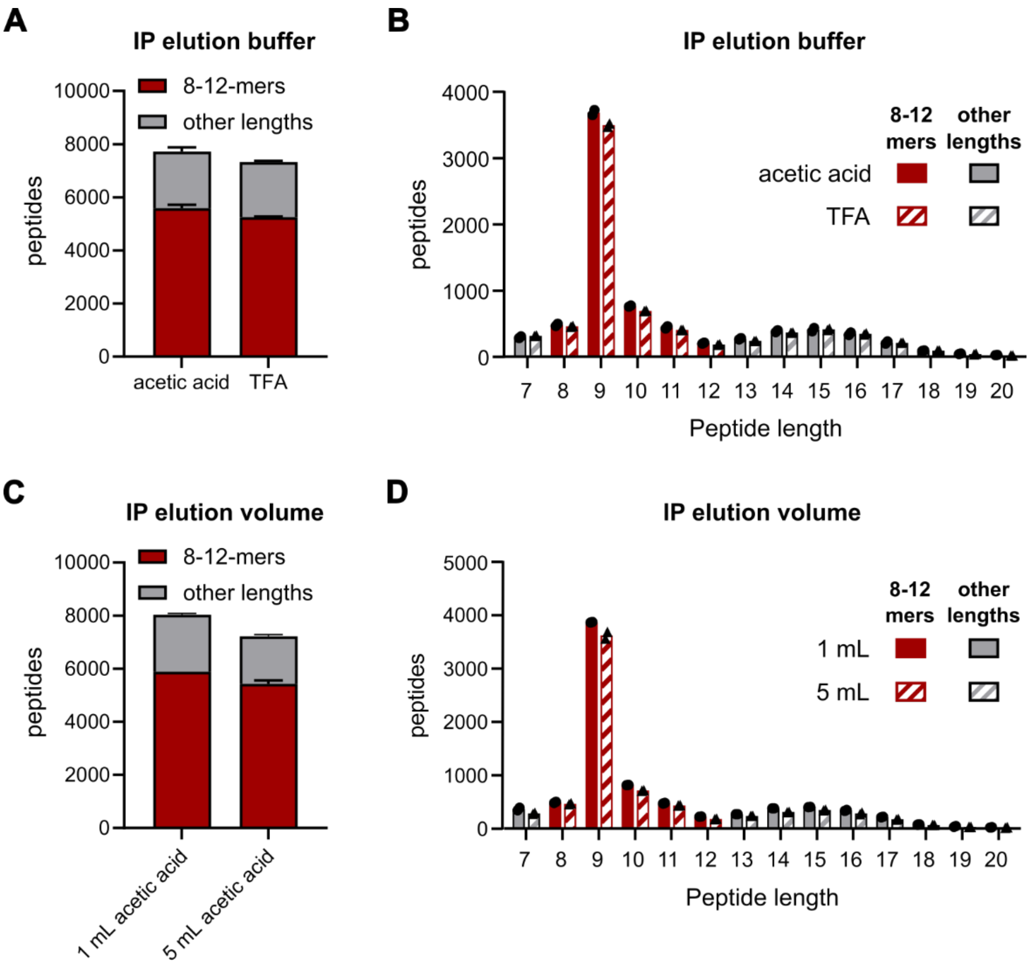


**Supplementary Figure S4. Optimization of the IP elution conditions.** Ten million JY cells were lysed in 100 µL lysis buffer and subjected to MHC-I IP using 300 µg W6/32 antibody crosslinked to 100 µg packed protein A beads. Peptide clean-up was performed according to the standard immunopeptidomics workflow, followed by sample injection on a timsTOF SCP instrument with the Evosep One chromatography system (25% sample injection). Data represent the average of 2 technical replicates. Peptides were identified using PEAKS (v11.0) (51). **(A, B)** Comparison of MHC-I IP elution using 5 mL 10% acetic acid or 5 mL 0.1% TFA. **(C, D)** Comparison of MHC-I IP elution using 5 mL or 1 mL 10% acetic acid. **(A, C)** Number of identified 8-12mer peptides (red) and peptides of other lengths (grey) identified in both conditions. **(B, D)** Peptide length distribution for both conditions.

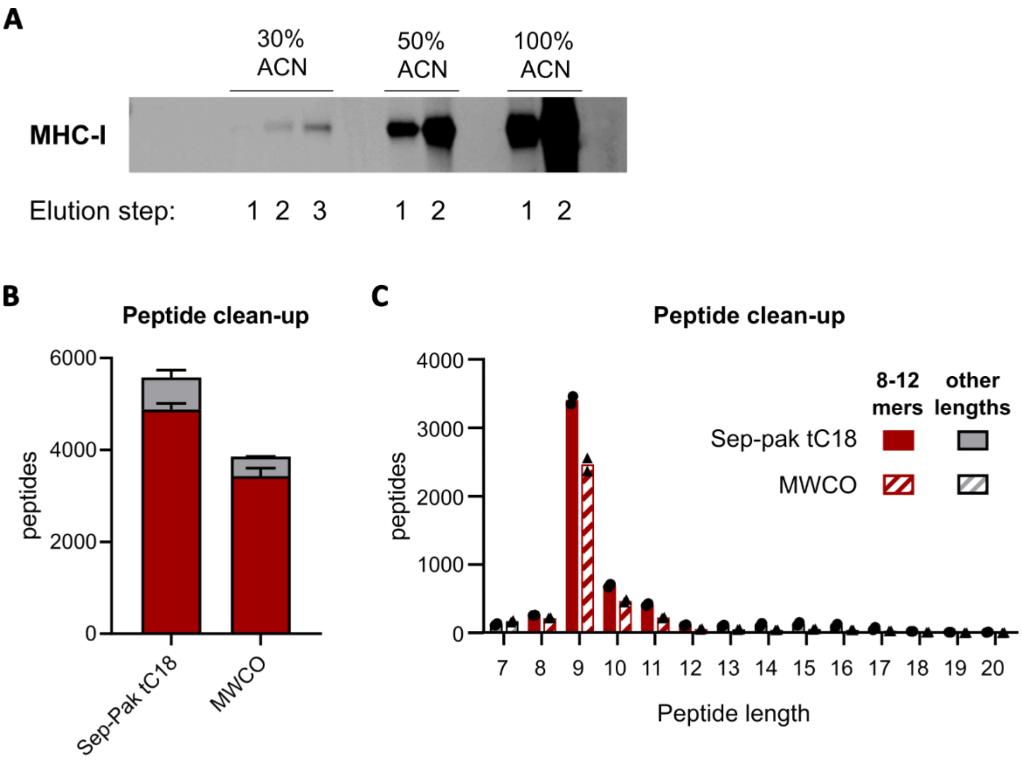


**Supplementary Figure S5**. **Optimization of the peptide purification method.** Ten million JY cells were lysed in 100 µL lysis buffer and subjected to MHC-I IP according to standard workflow. **(A)** Immunoblot analysis showing the elution of HLA class I proteins from Sep-Pak tC18 resin upon multiple elution steps in 0.1% TFA with increasing percentages of ACN. **(B-C)** Comparison of the number of identified peptides following clean-up using Sep-Pak tC18 96-well plates or 10 kDa molecular weight cut-off filters (MWCO). 10 million JY cells were lysed in 100 µL lysis buffer and subjected to MHC-I IP using 300 µg W6/32 antibody crosslinked to 100 µL packed protein A beads. Peptides were purified using either Sep-Pak tC18 96-well plates (100 mg) with elution in 25% ACN 0.1% TFA, or using a 10 kDa molecular weight cut-off filter followed by purification on a Sep-Pak tC18 96-well plate with elution in 40% ACN 0.1% TFA. Samples were run on the timsTOF SCP instrument with the Evosep One chromatography system (25% sample injection). Data represent the average of 2 technical replicates. Peptides were identified using PEAKS (v11.0) (51). **(B)** Number of identified 8-12mers (red) and peptides of other lengths per condition (grey). **(C)** Peptide length distribution for both conditions.

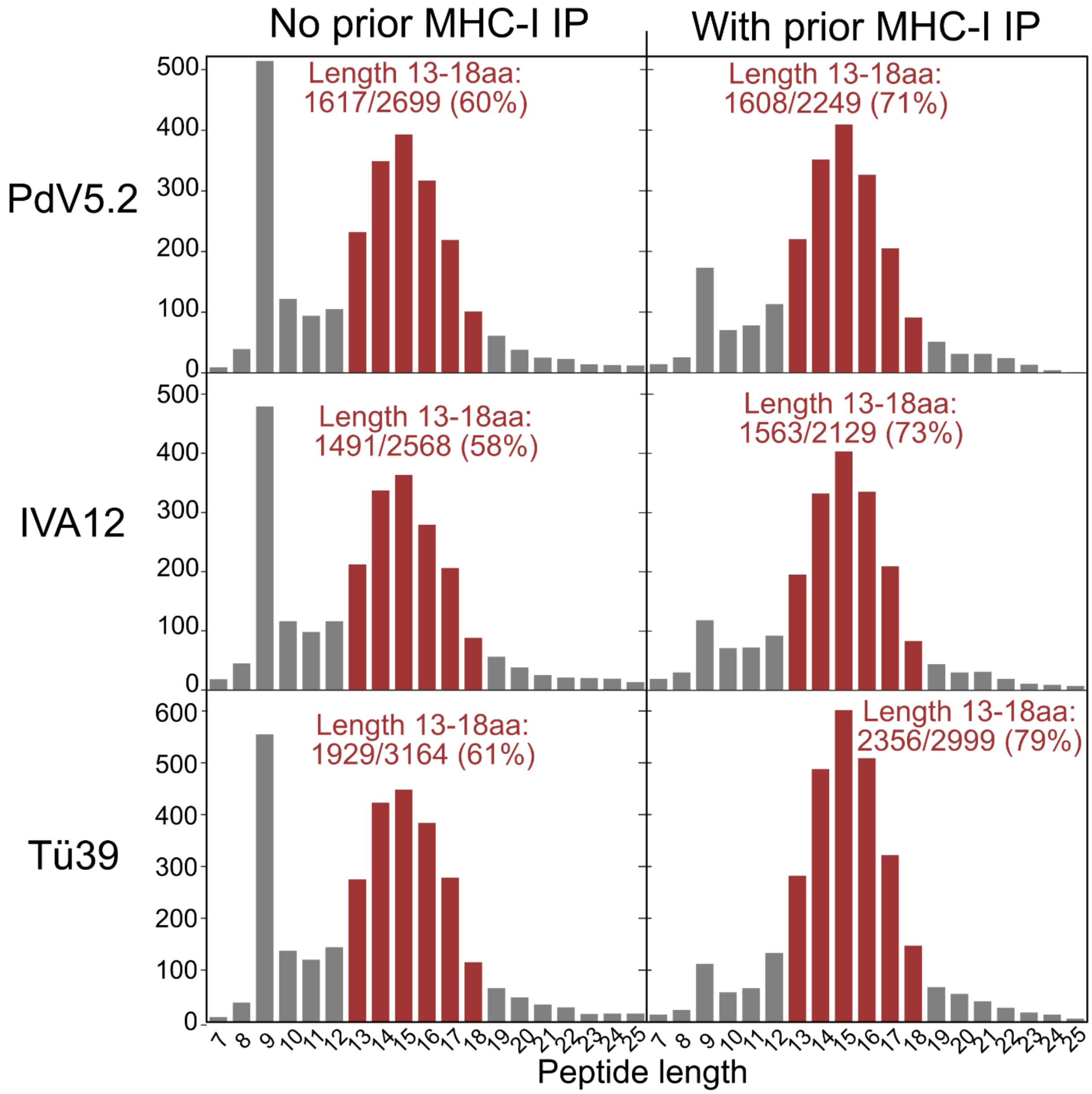


**Supplementary Figure S6. Identified peptide length histograms of MHC-II pulldowns testing different antibodies and without prior MHC-I pulldown.** Three pan MHCII antibodies were tested on the same input of JY cell material: PvD5.2, IVA12 and TU39 (*Top to down*). The number of unique identified peptide sequences was shown in function of peptide length in amino acids. Length 13 to 18 was indicated in red, representing a typical MHC-II compatible length peaking in MHC-II immunopeptidomics. Samples without a prior MHC-I pulldown (*Left*) and with a MHC-I pulldown (*Right*) were compared, the latter being a default approach to reduce the co-enrichment and detection of MHC-I peptides visible here by the 9-mer peak.

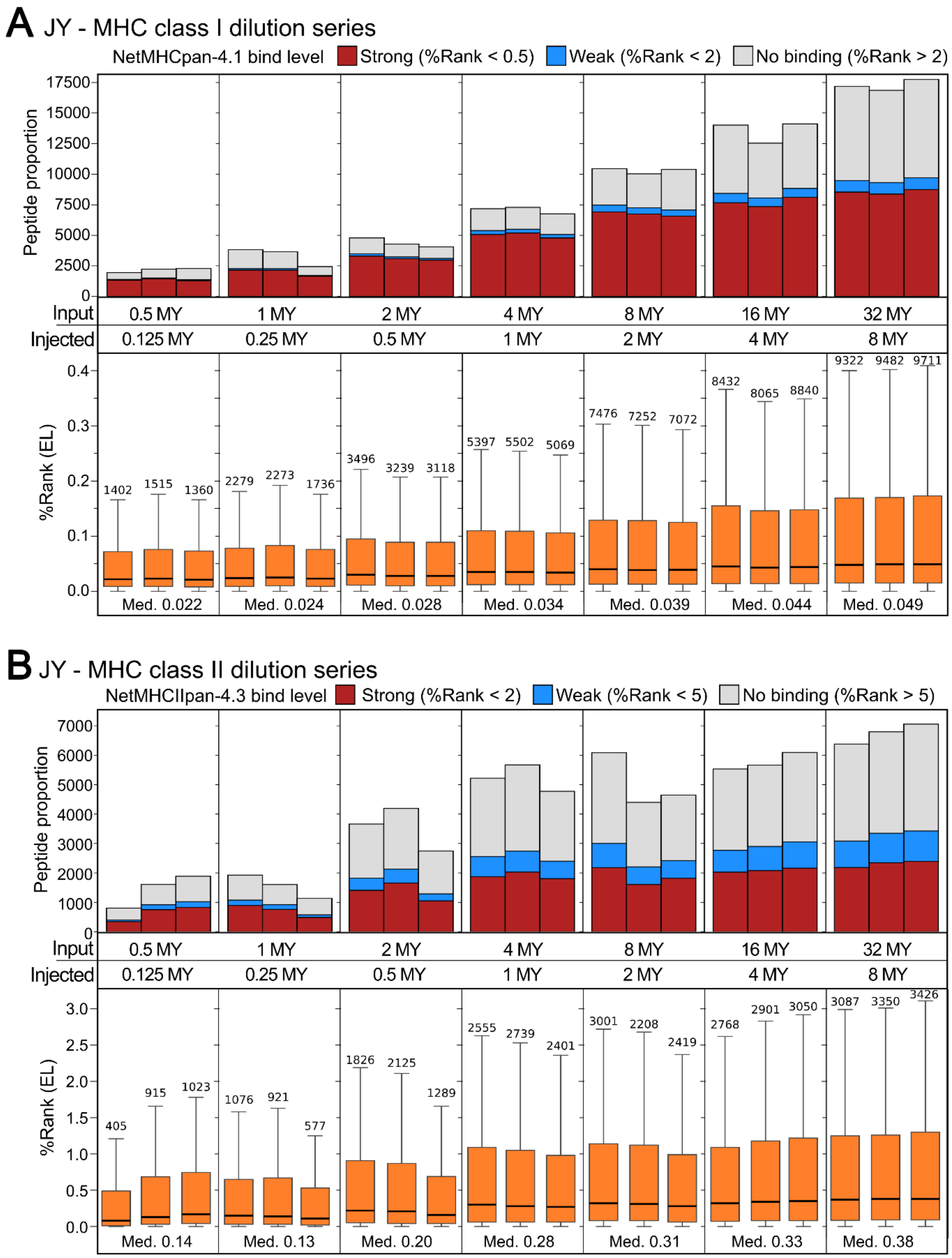


**Supplementary Figure S7. JY MHC class I and II predicted binding strengths. (A-B)** Identified peptides sequences per sample grouped per cell dilution series for MHC class I (**A**) and MHC class II (**B**) dilution series in JY cells. Each dilution was processed searched independently, and 25% of the input material was injected on timsTOF SCP. (*Top*) Peptides are colored according to predicted MHC binding strength by NetMHCpan-4.1 (53) for MHC class I (**A**) and NetMHCIIpan-4.3 (54) for MHC class II (**B**). (*Bottom*) Boxplot distributions of the %Rank score were shown for predicted binders in each individual sample. Per dilution, the median %Rank was displayed for all predicted MHC binders.

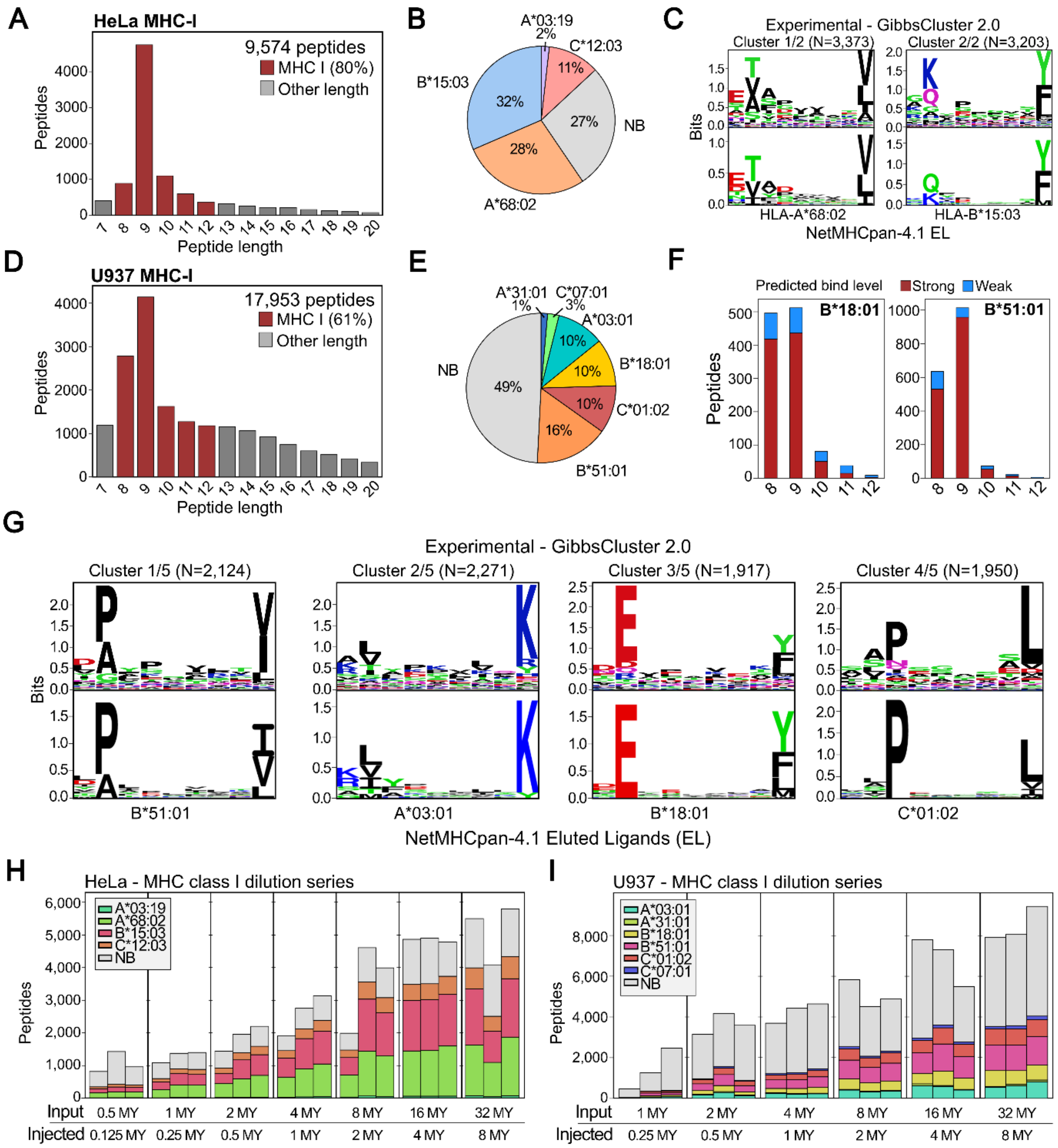


**Supplementary Figure S8. Immunopeptidomics on HeLa and U937 cell dilutions.** (**A, D**) Identified peptide length histogram of MHC class I pulldown in HeLa (**A**) and U937 (**D**). (**B, E**) Pie charts indicating the NetMHCpan-4.1 (53) predicted peptide binders to MHC alleles for HeLa (**B**) and U937 (**E**). Binders were defined as peptides with a %Rank < 2 and were assigned to the MHC allele with the lowest %Rank. Other peptides were classified as non-binders (NB, grey). (**C, G**) Unsupervised Gibbs clustering (55) of HeLa (**C**) and U937 (**G**) 8-12mer peptides reveals sequence logos match MHC class I allele eluted ligand (EL) motifs of netMHCpan-4.1 (53). (**F**) Number of predicted weak and strong peptides binders (%Rank <

2 and < 0.5, respectively) per peptide length to HLA-B18*01 and HLA-B51*01 in the U937 dataset. (**H-I**) Identified peptides sequences per sample grouped per cell dilution series for HeLa (**H**) and U937 (**I**). Each dilution was processed searched independently, and 25% of the input material was injected on timsTOF SCP. Peptides are colored according MHC binding prediction by netMHCpan-4.1.

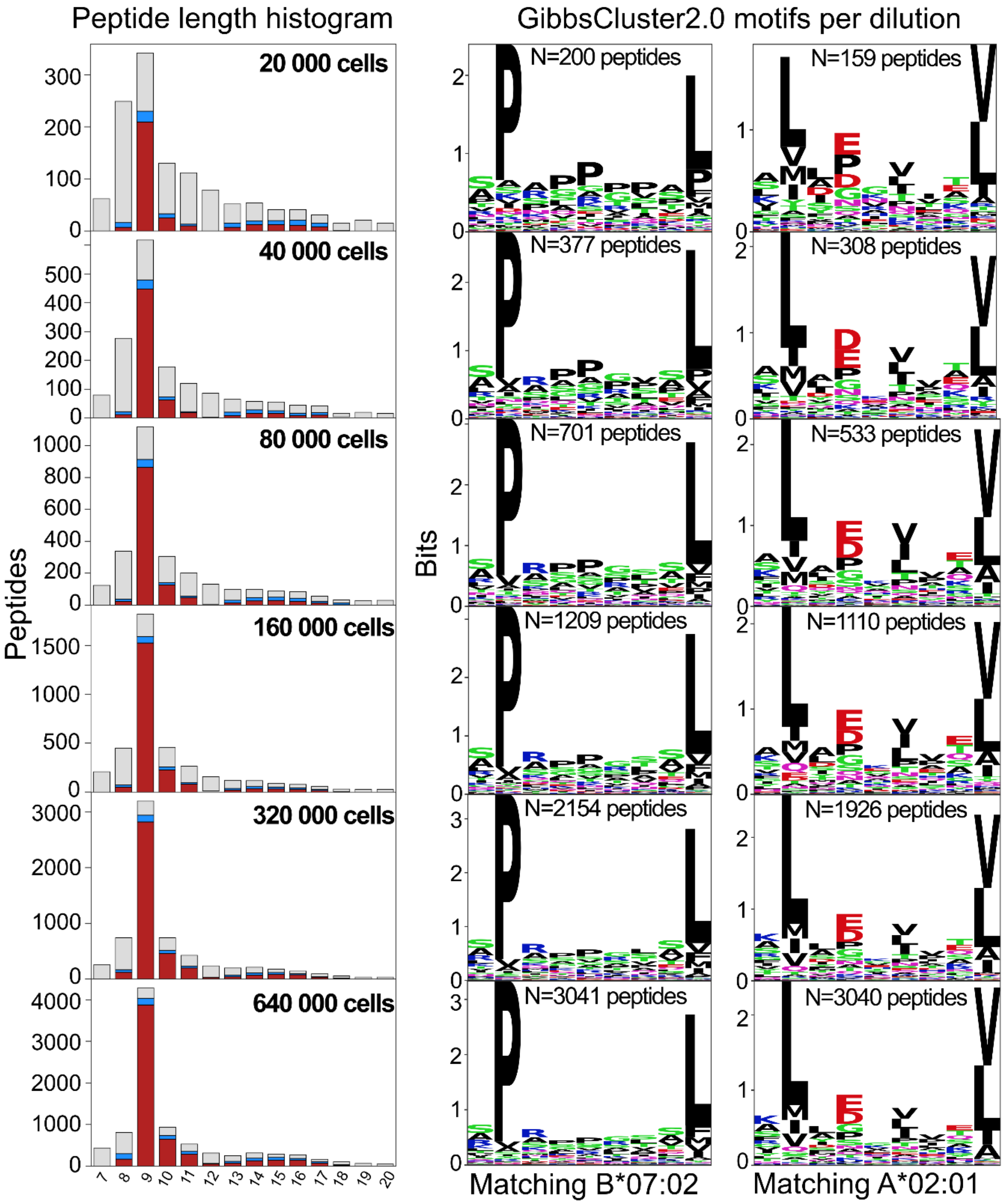


**Supplementary Figure S9. Peptide length and Gibbs clusters of ultrasensitive JY cell dilutions.** Peptide length distributions and sequence motifs corresponding to JY alleles HLA-A*02:01 and HLA-B*07:02 after unsupervised clustering identified 8-12mers by GibbsCluster2.0 (55) per JY cell dilution from 640,000 to 20,000 cells (*Bottom to top*). In the peptide length histograms, predicted weak and strong peptides binders (%Rank < 2 and < 0.5, respectively) were indicated in blue and red, respectively.

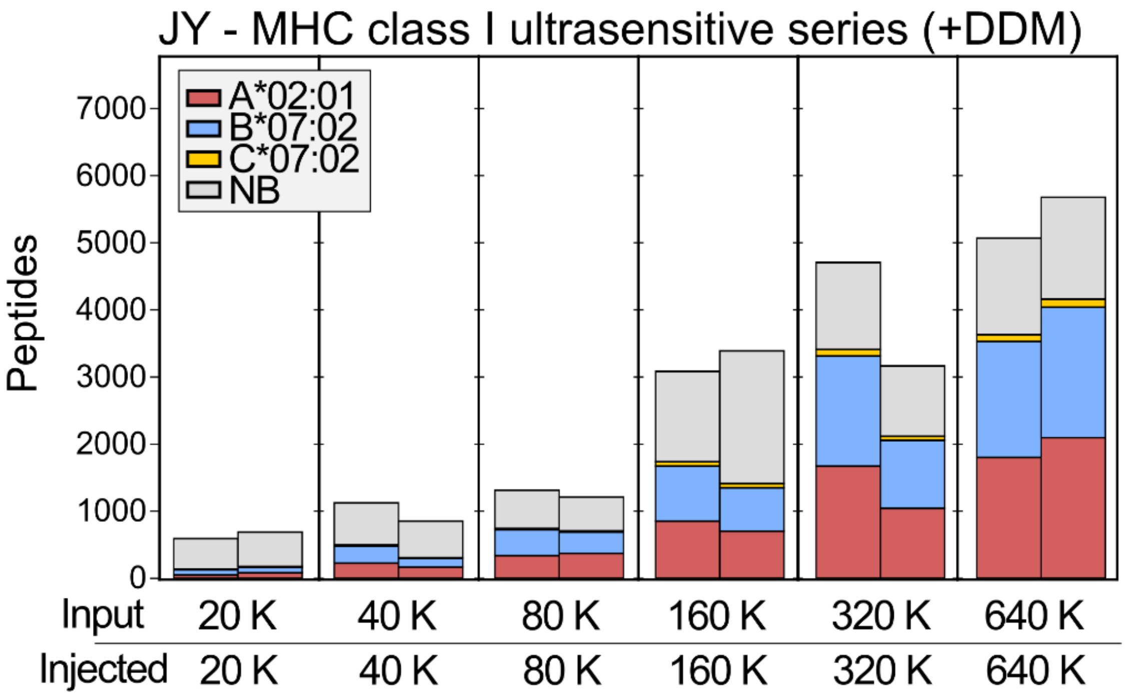


**Supplementary Figure S10. Ultrasensitive immunopeptidomics in presence of n-dodecyl-β-D-maltoside.** In an attempt to increase hydrophobic peptide recovery, a final volume of 0.02% n-dodecyl-β-D-maltoside (DDM) was incubated to each purified peptide mixture for 24 hours prior to LC-MS/MS injection. Identified peptides sequences per sample grouped per cell dilution series from 640,000 (640 K) down to 20,000 cells (20K). Each dilution was processed searched independently, and 100% of the input material was injected on timsTOF SCP. Peptides are colored according MHC binding prediction by netMHCpan-4.1 (53) for MHC class I JY alleles, indicating the predicted binders per MHC allele (%Rank < 2, lowest if multiple alleles).

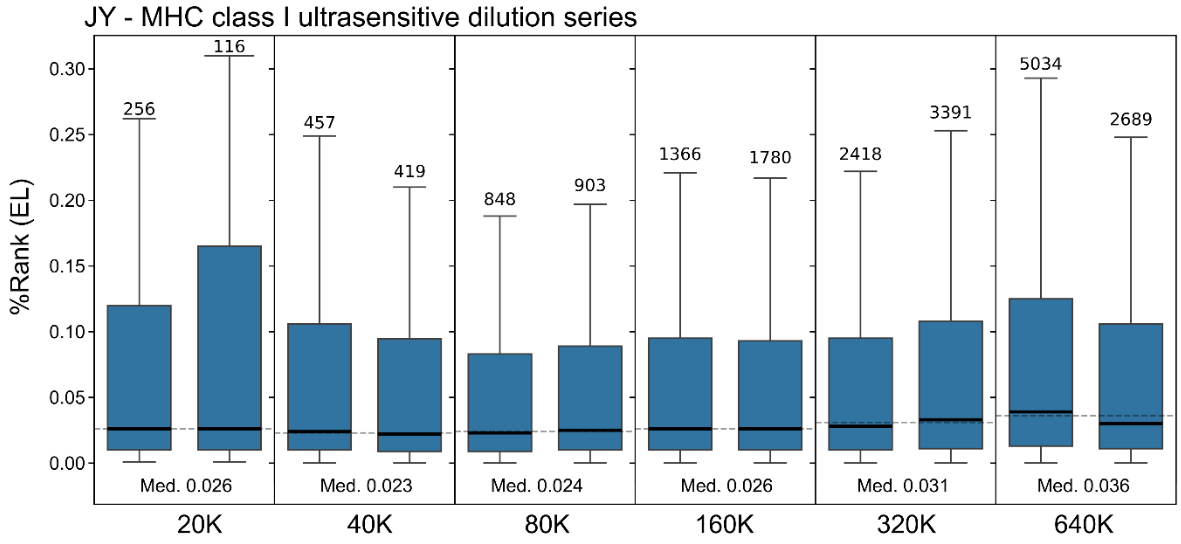


**Supplementary Figure S11. Predicted binding strength for identified immunopeptides in the ultrasensitive JY dilution series.** Boxplot distributions of the %Rank score calculated by NetMHCpan-4.1 (53) were shown for predicted binders in each individual sample. Per dilution, the median %Rank was displayed for all predicted MHC binders.

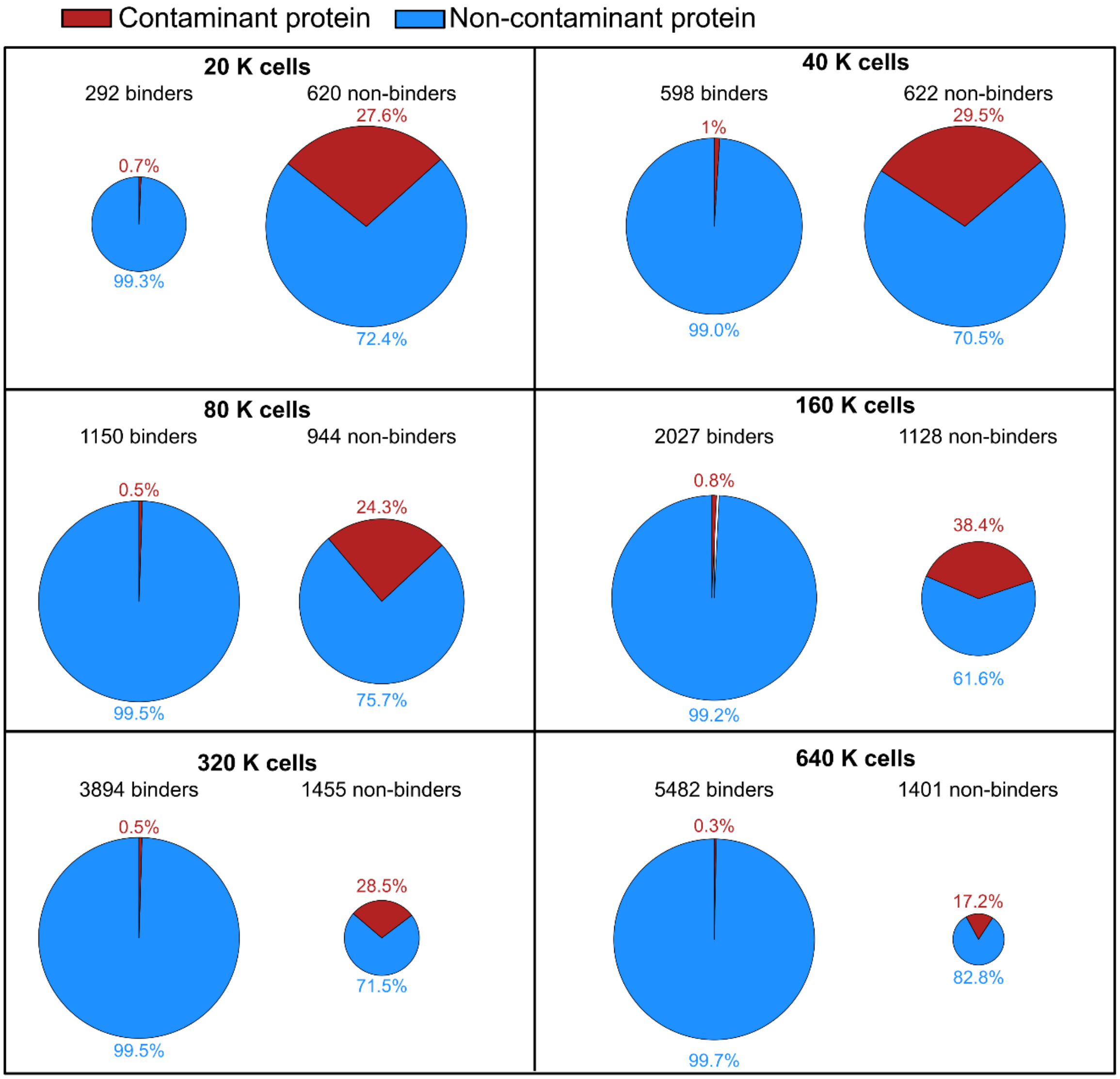


**Supplementary Figure S12. Predicted non-binders and contaminant proteins increase with decreasing cell amount inputs.** Per JY cell dilution the number of binders and non-binders was plotted in a proportional circle lay-out, indicating for each the number of peptides originating from non-contaminant proteins (blue) and those originating from the MaxQuant contaminants protein database (246 proteins).

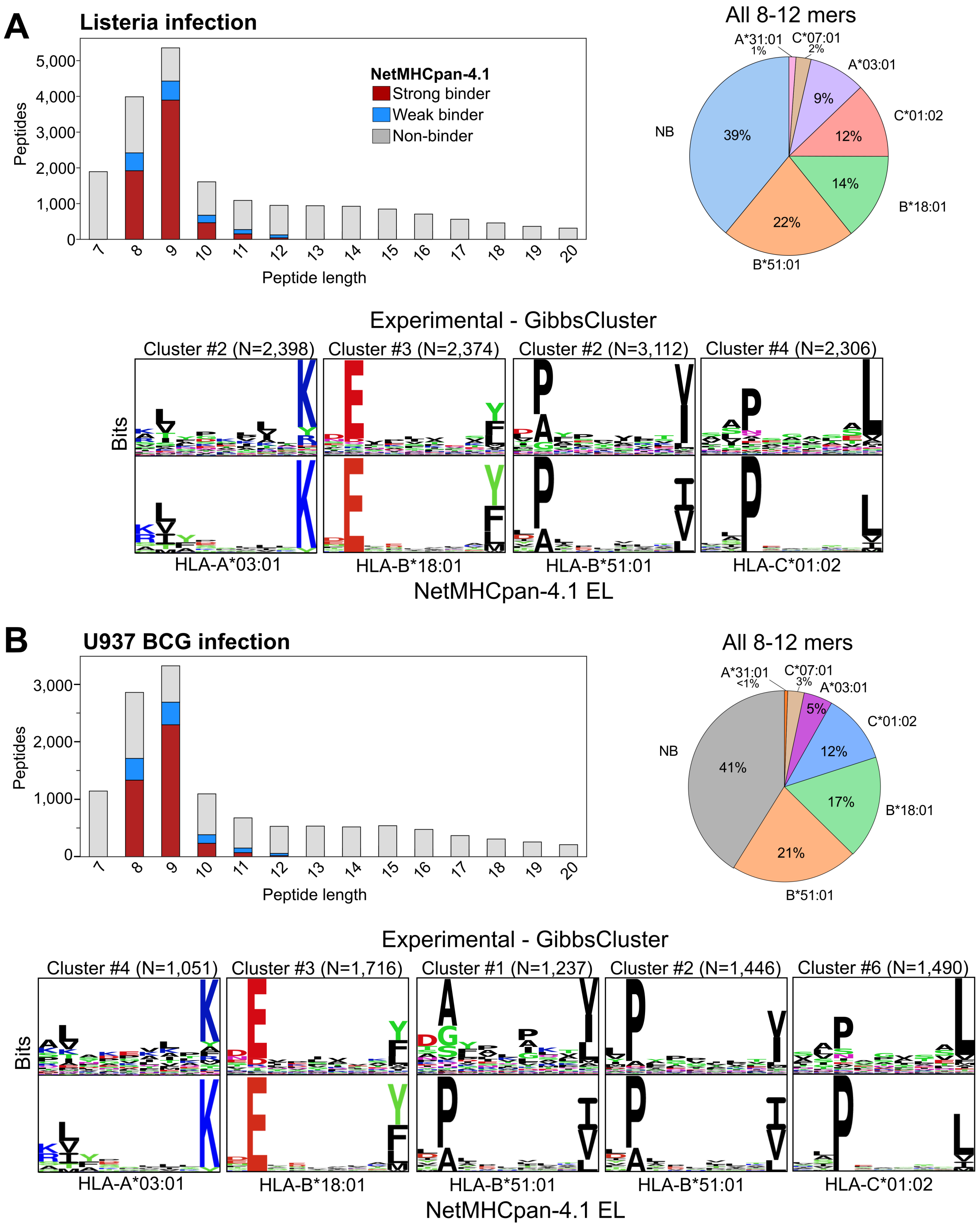


**Supplementary Figure S13. Immunopeptidomics quality control of U937 cell cultures infected by *Listeria monocytogenes* (A) and *Mycobacterium bovis* BCG (B).** A peptide length histogram showing the number of identified immunopeptides according their NetMHCpan-4.1 (53) binding prediction, with strong binders (%Rank < 0.5) in red, weak binders (%Rank < 2) in blue, and non-binders (%Rank > 2) in grey. For all 8-12mer peptides, the proportion of best-binding HLA alleles was given in a piechart. In addition, unsupervised GibbsCluster2.0 clustering (55) of 8-12mer peptides reveals sequence logos match MHC class I allele eluted ligand (EL) motifs of netMHCpan-4.1 (53).

**A** **Pairwise scatterplot**

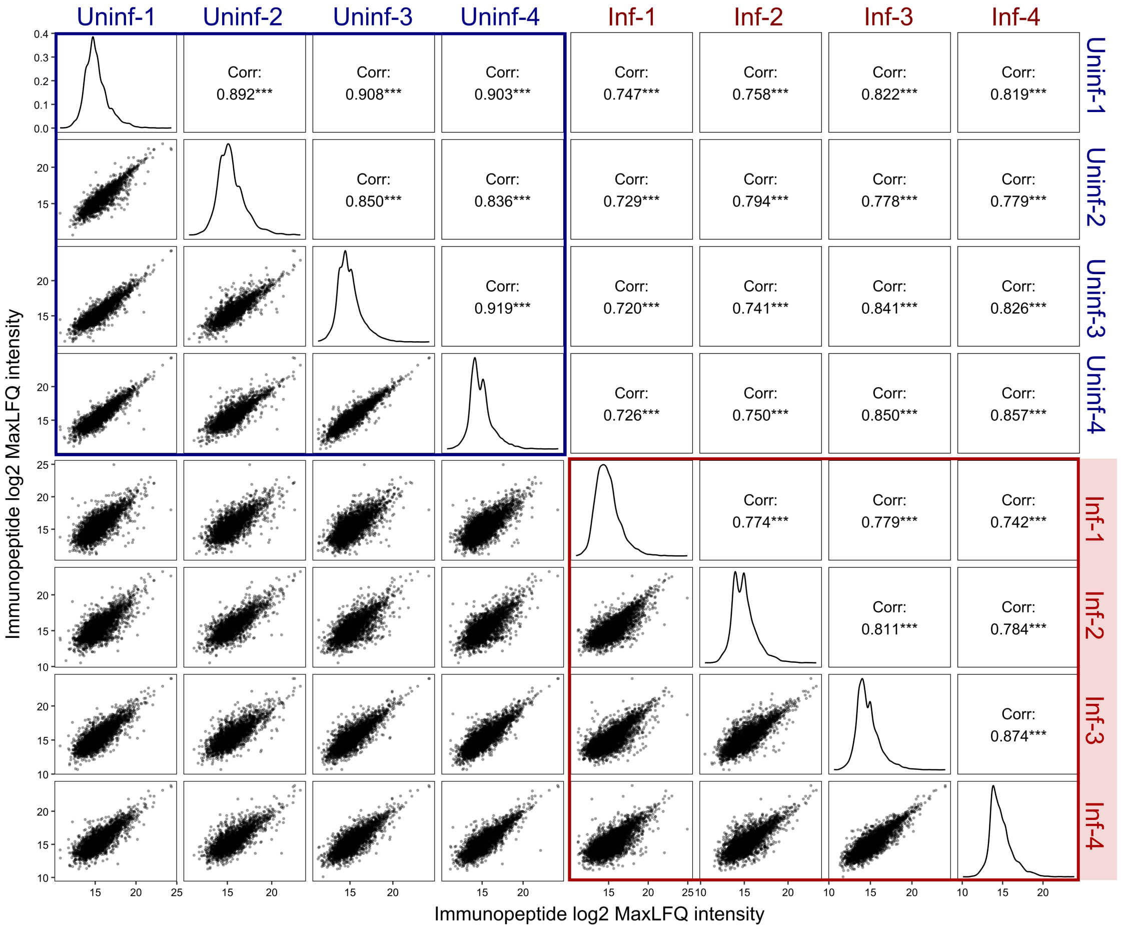


**B**

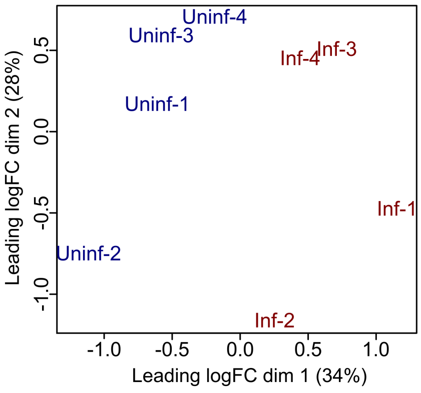


**Supplemental Figure S14. Quantitative reproducibility and variation between BCG-infected and uninfected samples.** (**A**) Pairwise scatter plots of log2-transformed MaxLFQ peptide intensities. The lower panels display pairwise scatter plots between samples, the upper panels show Pearson correlation coefficients, and the diagonal panels depict the distribution of intensities within each sample. Plots were made using the GGally (doi: 10.32614/CRAN.package.GGally) *ggpairs* function in R. Plots comparing uninfected ('Uninf') and BCG-infected ('Inf') samples were indicated in a blue and red rectangle, respectively. (**B**) Multidimensional scaling (MDS) plot of log2 normalized peptide intensities.

**Supplemental Methods**

This more detailed protocol explains the workflow for cross-linking beads and antibodies for immunoprecipitation (IP) of MHC-I and MHC-II complexes from different cell types, starting with 0.5 million cells. This protocol is also applicable to a variety of tissues.

Prepare all the buffers and solutions using ultrapure water and analytical-grade reagents. Rinse the glass bottles three times with ultrapure water before preparing the solutions. Always use freshly prepared solutions.

#### 1. Materials

##### *1.1. Antibody crosslinking beads*

1. Ultrapure water
2. Beads: Protein A-Sepharose® 4B Conjugate & Recombinant Protein G - Sepharose™ 4B
3. Tris-NaCl 50mM/150mM pH 8.
4. 0.2M sodium tetraborate decahydrate buffer pH 9.
5. Purified W6-32 anti-pan HLA-I antibody.
6. Purified Pdv5.2 anti-pan-HLA-II antibody.
7. Dimethyl pimelimidate (DMP)
8. 0.2M Ethanolamine solution pH 8: add 1.2 mL ethanolamine in 100 mL of Tris NaCl and adjust the pH by adding 37% HCl
9. Preservation buffer: PBS 1 with 0.02% NaN3.
10. Reusable Econo glass columns (#7374150, Bio-Rad) with a stand.
11. Gel electrophoresis system

##### *1.2. Purification of MHC-I and MHC-II binding peptides*

1. Cell pellets stored at -80 °C
2. Lysis buffer: 0.25% sodium deoxycholate (SDOC), 1% Octyl-beta D-glucopyranoside (OGP), 1 mM EDTA, 0.2 mM Iodoacetamide (IAA), 1 mM phenylmethylsulfonylfluoride, 1.25x complete™, Mini, EDTA-free Protease Inhibitor Cocktail, Tris-NaCl 50mM/150mM. Prepare fresh and keep on ice.
3. Tris-NaCl 50mM/150mM pH 8
4. 100% ACN
5. 0.1% TFA
6. 0.1% TFA with 25% ACN
7. 0.1% TFA with 40% CAN
8. 10% Acetic acid
9. Resolvex Tecan A200 positive pressure processor (Tecan) provides an automated processing solution for 96-well plates
10. Filter microplate, 96-well, polypropylene, with 0.7 µm glass fiber membrane, 2 mL/well, long drip (Agilent), part No. 201007-100
11. Sep-Pak tC18 96-well Plate(186002321), 100 mg Sorbent per Well, 37 - 55 µm (Waters)
12. 96-well “Collection” plates (2 mL per well, Waters)
13. 96-well “Waste” plates (2 mL per well, Waters)
14. 1.5 mL Eppendorf Safelock tubes

15. 1.5 mL protein LoBind® tubes(Eppendorf)
16. High-speed centrifuge
17. Vacuum centrifuge

*1.3. Mass spectrometry*

1. LC-MS/MS. We use a TimsTof SCP mass spectrometer online, coupled to a nano-LC system.

**2. Methods**

*2.1. Preparation of anti-MHC cross-linked beads for IP of MHC complexes*

1. Take two clean Econo columns labeled as “W6-32-protein A” and “PDV5.2-protein G”. Add 2 mL of fresh resuspended beads per column and allow the bead preservation buffer to drain off. Wash the beads 5 times with 5 mL of 50 mM/150 mM Tris-NaCl.
2. Load 3 mg of W6-32 and 1.5mg of PdV5.2 to the respective columns. (Make a 5 mL antibody solution by adding Tris-NaCl to load onto the beads. Take a 50 µL aliquot of antibody solution for QC silver staining.) Incubate for 1 h at room temperature (RT) on the tube roller at 80 rpm.
3. After incubation, collect the unbound antibody in a 15 mL Falcon tube. Take the 60 µL aliquot of unbound antibody solution for quality control.
4. Wash the columns with 5 mL of 0.2 M sodium tetraborate, pH 9, 5 times. Resuspend the beads with 5 mL of 0.2 M sodium tetraborate, pH 9. Transfer 50 µL of resuspended beads from each column to the Eppendorf tubes for QC to evaluate cross-linking efficiency.
5. Prepare the dimethyl pimelimidate (DMP) solution of 26 mg/5 mL of tetraborate and add it directly into the column. Close the column and incubate for 40-50 mins at RT on the tube roller at 80 rpm.
6. Remove the cap and allow the liquid to drain. Wash the beads 3 times with 0.2 M sodium tetraborate. Resuspend the beads in 5 mL of 0.2 M sodium tetraborate, and transfer 50 µL of the resuspended beads from each column to the Eppendorf tubes for QC.
7. Open the tip and let the liquid flow through. Wash the beads 3 times with 5 mL of 0.2 M ethanolamine solution. Close the column tips and add 5 mL of 0.2 M ethanolamine solution. Close the columns and rotate at RT for 2 h.
8. Open the column tips and caps. Allow the ethanolamine to flow through and wash the beads 5 times with 50 mM/150 mM Tris-NaCl. Close the column tip and resuspend the beads in 5 mL Tris-NaCl.Add storage buffer: Azide 0.02% (V/V). Close the tips and caps of columns tightly and store them at 4 °C for future use.

*2.2. Isolation of MHC-I and MHC-II peptides*

IP is performed in a semi-automated 96-well format using a positive-pressure processor (Tecan). The number of wells to be prepared for IP depends on the number of samples to be processed. The 96-well filter plates cannot be reused. Make sure to adjust the 96-well filter plates on the positive-pressure processor to achieve uniform pressure in each well and homogeneous solution flow.

*2.2.1. Preparation of 96-well plates for IP*

1. Label the 96-well filter plates “MHC-I plate” and “MHC-II plate”. Label two 96-well 2 mL plates as “Lysate collection” and “Elution collection”.
2. Mark the wells of the 96-well filter plate with the correct number and seal the remaining unused wells with the microplate sealing film.
3. Before loading beads, wash the marked wells using the processor: 1 mL 100% ACN twice, 1 mL 0.1% TFA twice, and 1 mL 50 mM/150 mM Tris-NaCl pH 8 three times.
4. Load 0.2 mL resuspended cross-linked beads per well “W6-32-ProteinA” and “PDV5.2-ProteinG” onto the “MHC-I plate” and “MHC-II plate” respectively.
5. Wash the beads five times, 1 mL 50 mM/150 mM Tris-NaCl pH 8, using the processor and push through the liquid by applying low-pressure (5-10%)

*2.2.2. Lysate preparation for the purification of MHC-I and MHC-II peptides*

1. Take out the cell pellets from -80 °C and thaw them on ice.
2. Add 0.1 mL cold lysis buffer to each cell pellet ranging from $0.5\times10^6$ to $32\times10^6$.
3. Lyse cells over 1 h by pipetting up and down 10X every 15 minutes.
4. Balance the centrifuge tubes and clear the lysate by centrifugation at 20,000 × g at 4°C for 10 min in a 1.5 mL Eppendorf tube to clear out big cell debris. Transfer the supernatant from each sample to the annotated tubes and centrifuge at 20,000 × g for 30 min to remove small cell debris. Transfer the clear lysate to the annotated tubes and keep them on ice. Take 3% of the lysate for quality control to run SDS gel.
5. For the infected cell lysate, the lysate is filtered out by using 0.22 µm Spin-X centrifuge tube filters at 16,000 × g.
6. For ultrasensitive workflow samples, lyse the cell pellets ranging from 20,000 to 640,000 in 50 µL of lysis buffer.

*2.2.3. IP of MHC-I and MHC-II*

1. Load 0.1 mL clear lysate from the samples into the assigned wells of the filter plate “MHC-I plate”. Stack the filter plate on top of the “Lysate collection” plate. Incubate the plates at + 4 °C for 1 h.
2. After incubation, place the “MHC-I plate” above the “Lysate collection” and push through the lysate by applying low pressure (course: 5%-10%) using the processor.
3. Load the collected lysate onto the beads into the assigned wells of the filter plate “MHC-II plate” and incubate the plate at 4 °C for 1 h.
4. Meanwhile, wash the “MHC-I plate” five times with 50 mM/150 mM Tris NaCl, pH 8, using the Tecan.
5. After incubation, place the “MHC-II plate” above the “Lysate collection” and push through the lysate by applying low pressure (course: 5%-10%) using the processor. Transfer the flow through the lysate to Eppendorf tubes, and take 3% of the lysate for quality control to run an SDS gel. Store the flow through lysate at -20 °C.
6. Wash the “MHC-II plate” five times, 50 mM/150 mM Tris NaCl, pH 8, using the processor.

*2.2.4. Purification of MHC-I and MHC-II-bound peptides*

1. Prepare two Sep-Pak tC-18 96-well plates. Label one as "MHC-I Sep-Pak" and the other as "MHC-II Sep-Pak". Activate the required wells of Sep-Pak plates twice 1 mL ACN and 3 times 1 mL miliQ 100%: 0.1% TFA. Drain the solutions by applying pressure.
2. Mount the "MHC-I plate" on top of the "MHC-I Sep-Pak" and the "MHC-II plate" on top of the "MHC-II Sep-Pak respectively. Elute the HLA-I and HLA-II filter plates with 0.2 mL 10% Acetic acid five times. Load 0.2 mL of Acetic acid each time, allow gravity flow for 5 minutes, then apply pressure.
3. Remove the "MHC-I plate" and "MHC-II plate" after acetic acid elution.
4. Wash the "MHC-I Sep-Pak" and "MHC-II Sep-Pak" plates three times with 1 mL of miliQ 100%, 0.1% TFA.
5. Place the "MHC-I Sep-Pak" plate on top of the "Elution collection-I" plate and Elute the MHC-I peptides with three times 500 µL 25% ACN.
6. Place the "MHC-II Sep-Pak" plate on top of the "Elution collection-II" plate and Elute the MHC-II peptides three times with 500 µL 40% ACN.
7. Transfer the elution containing peptides from the collection plates to the labeled protein LoBind tubes.
8. Dry the peptides completely using speed vacuum centrifugation. Store the dried peptides in a LoBind tube at -20°C until LC-MS analysis.
9. Resuspend the dried peptides in 50 µL of loading solvent A (0.1% TFA in water/acetonitrile (ACN) (99.5:0.5, v/v)). Vortex and centrifuge briefly. Sonicate the tubes for 5 min, then transfer the resuspended peptides to the correct labeled MS vials.
10. Place the MS vials in the autosampler to run on the mass spectrometer. TimsTOF SCP methods and following data analysis are described in the Experimental Procedures in the manuscript main text. Other instruments, acquisition methods and data analysis workflows can be used.
